# Huntington’s disease LIG1 modifier variant increases ligase fidelity and suppresses somatic CAG repeat expansion

**DOI:** 10.1101/2025.07.15.664798

**Authors:** Eunhye Lee, Wonju Kim, David H. Beier, Yejin Lee, Marina Kovalenko, Faaiza Saif, Esaria Oliver, Ryan Murtha, Marissa A. Andrew, Tammy Gillis, Brigitte Demelo, Bhairavi Srinageshwar, Jayla Ruliera, Diane Lucente, Seung Kwak, Ramee Lee, Ricardo Mouro Pinto, Marcy E. MacDonald, James F. Gusella, Patrick J. O’Brien, Vanessa C. Wheeler, Ihn Sik Seong

## Abstract

Huntington’s disease (HD) is a fatal neurodegenerative disorder caused by inheriting an expanded CAG repeat tract in the huntingtin gene (*HTT*) that further expands in somatic cells over an individual’s lifetime. Genome-wide association studies have provided critical insight into factors that modify the course of disease. These include DNA repair genes that alter the rate of somatic expansion and other genes that do not appear to directly influence this process. One modifier gene is DNA ligase 1 (*LIG1*), in which a variant specifying a lysine to asparagine substitution (K845N) is associated with a profound (7-8 year) delay in the onset of motor signs. Here, we have taken a multifaceted approach to gain insight into the protective nature of this variant in HD. We demonstrate using *in vitro* ligase assays and enzyme kinetics that K845N enhances discrimination towards mismatched substrates and increases repair fidelity. Consistent with increased ligation fidelity, K845N confers protection against oxidative stress in cell-based assays. Finally, we demonstrate that the mouse LIG1 K843N orthologue suppresses somatic CAG expansion in HD knock-in mice. Overall, our data provide evidence that altered LIG1 function due to the K845N substitution may contribute to HD clinical delay by slowing somatic expansion in the brain and protecting the genome globally against damage. Significantly, our results provide a mechanistic foundation for considering DNA ligase fidelity as a therapeutic target in HD and potentially in other trinucleotide repeat disorders.

**Significance Statement:** We analyzed a missense variant in DNA Ligase 1 (K845N) that is associated with a profound delay in the onset of Huntington’s disease (HD). We find that K845N enhances substrate discrimination towards mismatched substrates, thus increasing repair fidelity, conferring protection against oxidative stress and slows somatic expansion of the HD CAG repeat. Our observations provide insight into underlying mechanisms of disease modification and suggest avenues that can be harnessed for disease-modifying therapeutic intervention.

**Classification:** Biological Sciences, Genetics

## Introduction

Huntington’s disease (HD) (MIM #143100) is a progressive, autosomal dominant neurodegenerative disorder characterized by motor dysfunction, cognitive decline, and behavioral disturbances (1). It is caused by inheriting an expanded CAG trinucleotide repeat (≥ 36 repeat units) within the first exon of the huntingtin gene (*HTT*) located on chromosome 4 (2). The expanded CAG repeat is unstable in somatic cells and progressively increases in length over the life of HD individuals, particularly in neurons in the brain (3–8). The length of the inherited CAG repeat is the most significant genetic factor influencing the age at motor onset (AAO) of HD, with longer repeat lengths associated, on average, with an earlier onset of motor symptoms (9–11). While repeat length accounts for a substantial portion (∼60%) of the variation in AAO among individuals with HD, there remains considerable variability in AAO (± ∼20 years) that is unexplained by inherited repeat length and which is heritable, indicating additional genetic factors that contribute to disease (10).

Identifying these modifying factors is essential to gain a comprehensive understanding of HD pathogenesis. Insights into these modifiers may improve predictions of the onset and progression of HD and inform the development of novel disease-modifying therapies. Genome-wide association studies (GWAS) have emerged as a powerful tool for systematically scanning the genomes of large cohorts of HD patients, enabling the identification of genetic variants associated with specific disease characteristics, such as AAO (12–17). Notably, these GWAS have identified genes involved in DNA repair pathways, in particular in mismatch repair (MMR), as modifiers of HD AAO and other clinical measures, and together with single cell-based and mechanistic studies, support a critical role for the somatic expansion of the CAG repeat in determining the timing of disease phenotypes (8, 18). Other modifier genes do not directly implicate somatic CAG expansion and may be involved in neuronal vulnerability/toxicity mechanisms (15, 17).

One of the genetic modifiers, associated with modification of multiple HD clinical landmarks, is *LIG1* (chromosome 19q13.33), encoding DNA ligase 1 (LIG1). The most recent GWAS (17) defined two distinguishable *LIG1* modifier effects, named 19AM1 and 19AM3. The top single nucleotide variant (SNV), whose minor allele tags the 19AM3 modifier effect is rs145821638 (GRCh38 - Chr19:48117686-C-A), specifies a lysine to asparagine substitution in LIG1, at position 845 (K845N). This uncommon variant (minor allele frequency 0.18 in Europeans, ∼0.14 across all gnomAD v4.1.0 populations) is associated with a 7-8-year delay in AAO, standing out as one of the strongest HD modifier effects detected to date. There are no other variants of comparable effect size and significance on the phased haplotype of the *LIG1* region from 19AM3 modifier chromosomes, arguing that K845N is the source of the modifier effect. The profound impact associated with this variant provides a strong rationale for understanding the mechanism of disease modification, and has potential to translate this knowledge into a therapeutic delay of disease onset.

LIG1 is an essential enzyme in mammalian development (19, 20) that catalyzes the formation of phosphodiester bonds to seal single-strand breaks in DNA. LIG1 plays a crucial role in completing DNA replication, where it joins Okazaki fragments on the lagging strand, and in DNA repair pathways, including base excision repair (BER), nucleotide excision repair (NER), and MMR (21, 22). High fidelity ligation by LIG1 is essential for maintaining the integrity of the genome by preventing the loss of genetic information and harmful mutations. Errors in ligation, particularly those involving mismatched bases, can lead to genome instability, which has been implicated in various diseases, including cancer and immunodeficiency (22).

LIG1 catalyzes the DNA ligation reaction through a three-step mechanism (23). In the first step, adenylylation, LIG1 reacts with ATP to covalently attach an adenosine monophosphate (AMP) moiety to a specific lysine residue in its active site (K568), forming a LIG1-AMP intermediate. The second step, adenylyl transfer, involves the transfer of AMP from LIG1 to the 5’ phosphate end of the DNA nick, resulting in an AMP-DNA intermediate. Finally, in the nick sealing step, the 3’ hydroxyl group at the DNA nick attacks the 5’ phosphodiester bond, displacing AMP and sealing the DNA backbone. The high fidelity of LIG1 relies on the ability of its active site residues to recognize and properly align the DNA ends, ensuring efficient and accurate ligation (24, 25). Some biallelic missense mutations in *LIG1* cause a recessive primary immune deficiency referred to as LIG1 Syndrome or IMD96 (MIM #619774) (26–28) associated with reduced LIG1 catalytic activity and altered DNA binding, highlighting potential for specific amino acid substitutions to alter LIG1 function. No such LIG1 Syndrome-associated mutations were identified in HD modifier GWAS cohorts, presumably due to their very low population frequencies (Table 1). However, homozygotes for the A allele specifying LIG1 K845N are observed in the population (gnomADv4.1.0) but this variant has not been reported to cause LIG1 Syndrome, hinting that any impact of the K845N change on LIG1 activity is distinct from the effects causing LIG1 Syndrome. Indeed, such an impact may not be disease-causing *per se* but rather become evident only under particular circumstances, such as an intersection with the HD pathogenic mechanism.

**Table 1.**
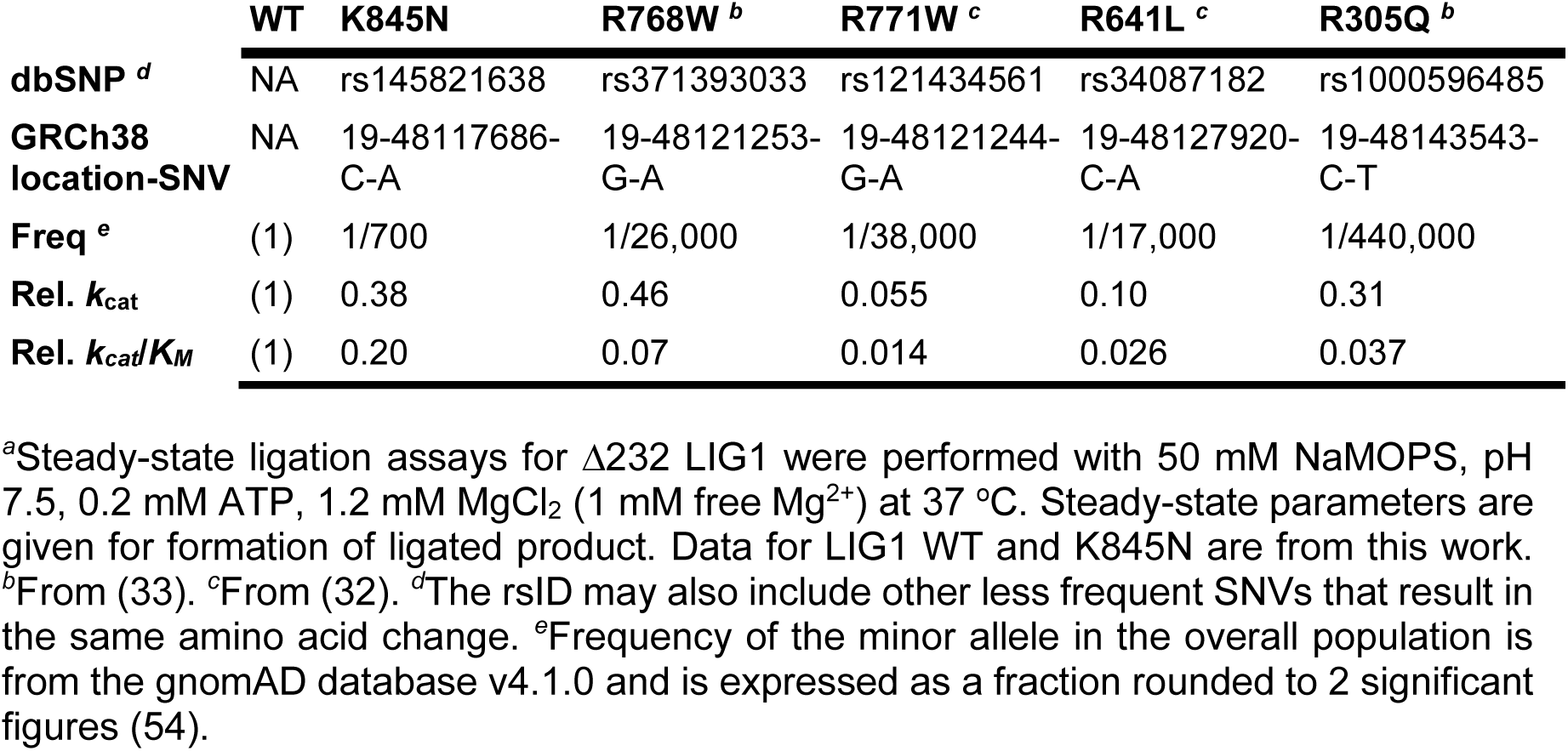
Minor allele frequency and relative ligase activity for LIG1 variants *^a^*

|  | WT | K845N | R768W <sup>b</sup> | R771W <sup>c</sup> | R641L <sup>c</sup> | R305Q <sup>b</sup> |
| --- | --- | --- | --- | --- | --- | --- |
| **dbSNP <sup>d</sup>** | NA | rs145821638 | rs371393033 | rs121434561 | rs34087182 | rs1000596485 |
| **GRCh38 location-SNV** | NA | 19-48117686-C-A | 19-48121253-G-A | 19-48121244-G-A | 19-48127920-C-A | 19-48143543-C-T |
| **Freq <sup>e</sup>** | (1) | 1/700 | 1/26,000 | 1/38,000 | 1/17,000 | 1/440,000 |
| **Rel. $k_{cat}$** | (1) | 0.38 | 0.46 | 0.055 | 0.10 | 0.31 |
| **Rel. $k_{cat}/K_M$** | (1) | 0.20 | 0.07 | 0.014 | 0.026 | 0.037 |

<sup>a</sup>Steady-state ligation assays for $\Delta 232$ LIG1 were performed with 50 mM NaMOPS, pH 7.5, 0.2 mM ATP, 1.2 mM MgCl<sub>2</sub> (1 mM free Mg<sup>2+</sup>) at 37 °C. Steady-state parameters are given for formation of ligated product. Data for LIG1 WT and K845N are from this work. <sup>b</sup>From (33). <sup>c</sup>From (32). <sup>d</sup>The rsID may also include other less frequent SNVs that result in the same amino acid change. <sup>e</sup>Frequency of the minor allele in the overall population is from the gnomAD database v4.1.0 and is expressed as a fraction rounded to 2 significant figures (54).

The biochemical properties of the LIG1 K845N variant and its potential as a contributor to a mechanism that delays HD pathogenesis are unknown. In this study, we have taken a multifaceted approach to this question that includes *in vitro* biochemical, cell-based and mouse studies. Our results demonstrate that LIG1 K845N reduces ligase activity towards mismatched substrates, with kinetic studies demonstrating increased ligation fidelity. In cell-based models of oxidative damage, we show protection by the K845N variant against cellular toxicity, consistent with a reduction in ligating pro-mutagenic nicks. Finally, we demonstrate that the mouse ortholog of the human LIG1 K845N variant, LIG1 K843N, suppresses somatic CAG expansion in HD knock-in mice. Taken together, these comprehensive approaches provide novel insight into the role of the LIG1 K845N variant in maintaining genomic integrity, implicate mechanisms of action of this modifier variant and support further investigation to harness insights into K845N for modifying disease outcomes in HD.

## Results

### LIG1 K845N shows reduced ligase activity *in vitro* that is sensitive to the ligation context, particularly pro-mutagenic 3’ mismatches

The K845N variant of LIG1 substitutes a positively charged lysine with a polar asparagine within the conserved oligonucleotide binding-fold domain (OBD) that is essential for DNA binding and positioning (Fig. 1*A* and *B*). K845 participates in interdomain interactions between the OBD and the adenylylation domain (AdD) (*SI Appendix*, Fig. S1). To investigate the impact of the K845N substitution on DNA ligase activity, we assessed the *in vitro* ligase activity of LIG1 with lysine at position 845 (LIG1 WT) and LIG1 with asparagine at position 845 (LIG1 K845N) using acrylamide gel-based analyses (Fig. 1*C*). Both full-length LIG1 WT and K845N proteins were purified in their adenylylated forms which allowed us to measure the concentration of active enzyme using single turnover ligation in the absence of ATP (*i.e*., active site titrations; *SI Appendix*, Fig. S2). LIG1 WT and K845N showed similar active concentrations of 89% and 92%, demonstrating that the K845N substitution does not impact the purification of the adenylylated enzyme.

**Figure 1.**
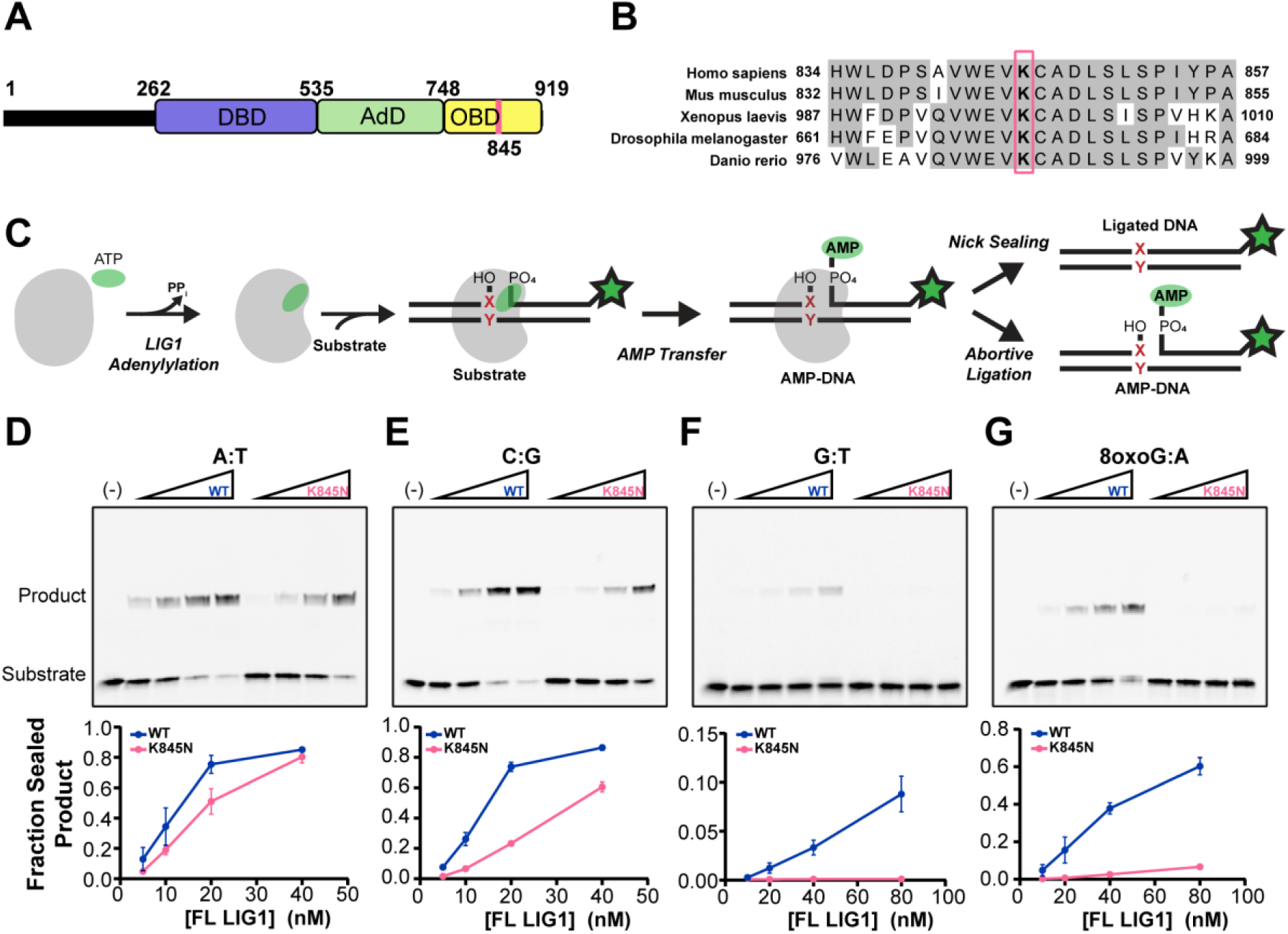
LIG1 K845N shows reduced ligase activity towards mismatched substrates. (A) Domain architecture of LIG1. The DNA binding domain (DBD), adenylylation domain (AdD), and OB-fold domain (OBD) comprise the catalytic core. (B) Multiple sequence alignment of LIG1 orthologs by Clustal Omega. The position of K845 is indicated by the red box. (C) Illustration of LIG1 reaction. LIG1 is first adenylylated by ATP at K568, forming a LIG1-AMP intermediate. The second step, adenyl transfer, involves the transfer of AMP from LIG1 to the 5’ phosphate end of the DNA nick, resulting in a DNA-AMP intermediate. Finally, in the nick sealing step, the 3’ hydroxyl group at the DNA nick attacks the 5’ phosphodiester bond, displacing AMP and sealing the DNA backbone. LIG1 may also release premature AMP-DNA intermediates, known as abortive ligation. Our *in vitro* ligation reaction uses nicked substrate generated by annealing three oligonucleotides, one of which is labeled with FAM (green star), allowing substrate, aborted DNA-AMP and ligated product to be distinguished. X:Y = canonical or mismatched base pair. (D-G) Representative gels and quantified fraction of ligation product showing the ligase activity of full-length LIG1 WT and K845N in the context of nucleotide pairs at the 3’ nick position. A:T (D), C:G (E), G:T (F), 8-oxoG:A (G) containing nicked 34mer DNA (300 nM) was incubated with increasing concentrations of LIG1 proteins for 5 min at 37°C. Reactions were performed with 1 mM ATP, 10 mM MgCl_2_, 50 mM MOPS pH 7.5 and 150 mM NaCl. Reactants were analyzed on 15 % TBE-urea polyacrylamide mini gels (note that in these gels substrate and AMP-DNA are not resolved). The line graphs show the quantification of the fraction of ligated product from three independent experiments (Mean ± SD).

We compared ligase activity of full-length LIG1 K845N to WT in a standard ligase assay buffer containing a saturating amount of Mg^2+^ (∼10 mM) and varying concentrations of enzyme (Fig. 1*D – G*). We used 34mer canonical nicked DNA substrates with different nucleotides at the 3’ end of the nick (Fig. 1*C*; *SI Appendix*, Table S1). Ligase activity was quantified as the ratio of the intensity of the ligated DNA product relative to the summed intensities of the ligated DNA, aborted AMP-DNA, and unreacted substrate. Using a dA:dT (hereafter simplified to A:T) substrate, we observed very similar ligase activity for LIG1 K845N and WT (Fig. 1*D*). In contrast, there was a modest, but significant decrease in ligase activity for K845N when a 3’ C:G substrate was tested (Fig. 1*E*). This pattern was also observed with T:A and G:C substrates, where complementary base pairing was reversed (*SI Appendix*, Fig. S3). From these data it is apparent that K845N is highly active as a DNA ligase, however it shows context-dependent effects that differ from those of LIG1 WT.

LIG1-catalyzed nick sealing completes DNA replication and repair pathways, and therefore the ability to distinguish mismatches from correctly paired bases helps to maintain genomic integrity. We next tested the pro-mutagenic contexts that would result from polymerase incorporation errors using all 12 possible mismatches at the 3’-OH side of the nicked DNA (Fig. 1*F* and *SI Appendix*, Figs. S4 & S5). Under our experimental conditions, the K845N variant showed no detectable ligation activity on any of the mismatched substrates, whereas LIG1 WT exhibited weak but detectable activity on the G:T, C:T, T:C, T:G, C:A, and A:C mismatches (Fig. 1*F* *and SI Appendix, Fig. S4*). For the remaining six base pair combinations, WT showed little or no ligase activity and K845N also showed no detectable ligase activity (*SI Appendix*, Fig. S5). This systematic screen of mismatched ligation contexts demonstrated that K845N has reduced activity toward sealing a variety of pro-mutagenic nicks.

LIG1 is known to be challenged in distinguishing an 8-oxoG:A mispair relative to other types of mismatches due to its stable Hoogsteen base pairing (25, 29–31) and yet this is a common pro-mutagenic substrate formed under oxidative conditions. We therefore compared the activity of LIG1 K845N and WT in the 3’ 8-oxoG:A ligation context. As was observed for the non-oxidized mismatches, K845N showed reduced ligase activity with 8-oxoG:A compared to LIG1 WT (Fig. 1*G*). Reduced ligation activity towards replication errors such as mismatches or incorporation of 8-oxo-dGTP from the oxidized nucleotide pool could elevate the overall fidelity of genome maintenance and support a protective function of K845N.

### K845N selectively aborts ligation of an 8-oxoG:A mismatch to enhance the fidelity of ligation

To elucidate how the K845N substitution alters the biochemical properties of LIG1, we performed a thorough kinetic characterization of Δ232 LIG1 under the same conditions that have previously been used to study other rare variants of LIG1 (26, 32, 33). This truncation removes the disordered and poorly conserved N-terminus of LIG1 which contains the nuclear localization sequence (24, 34). Active site titrations were used to determine the concentration of active Δ232 LIG1 (*SI Appendix*, Fig. S6). Representative time courses are shown in Fig. 2*A* comparing LIG1 WT and K845N with an undamaged nicked DNA substrate (C:G) and a substrate containing an 8-oxoG:A at the 3’ end of the nick. Similar to what was observed for the full-length LIG1 at higher Mg^2+^ concentration with a different sequence context (Fig. 1 *E* and *G*), K845N exhibited reduced activity on the undamaged substrate and substantially less activity on the 8-oxoG:A substrate. Using high resolution sequencing gels (denaturing PAGE), the abortive AMP-DNA species that is particularly prevalent in reaction of the 8-oxoG:A substrate is resolved from unreacted substrate. We next measured the initial rates of ligation at saturating concentration of DNA (1000 nM). Representative initial rates are shown for the canonical substrate in Fig. 2*B*. These data show that the reduced production of sealed DNA product in reactions catalyzed by K845N (solid lines) is accompanied by a detectable increase in the amount of AMP-DNA release (dashed lines; Fig. 2*B*). Analogous reactions performed with the 8-oxoG:A substrate show that AMP-DNA release by LIG1 WT is roughly on par with the amount of ligated DNA, whereas LIG1 K845N releases mostly AMP-DNA and fails to complete the ligation reaction (Fig. 2*C*).

**Figure 2.**
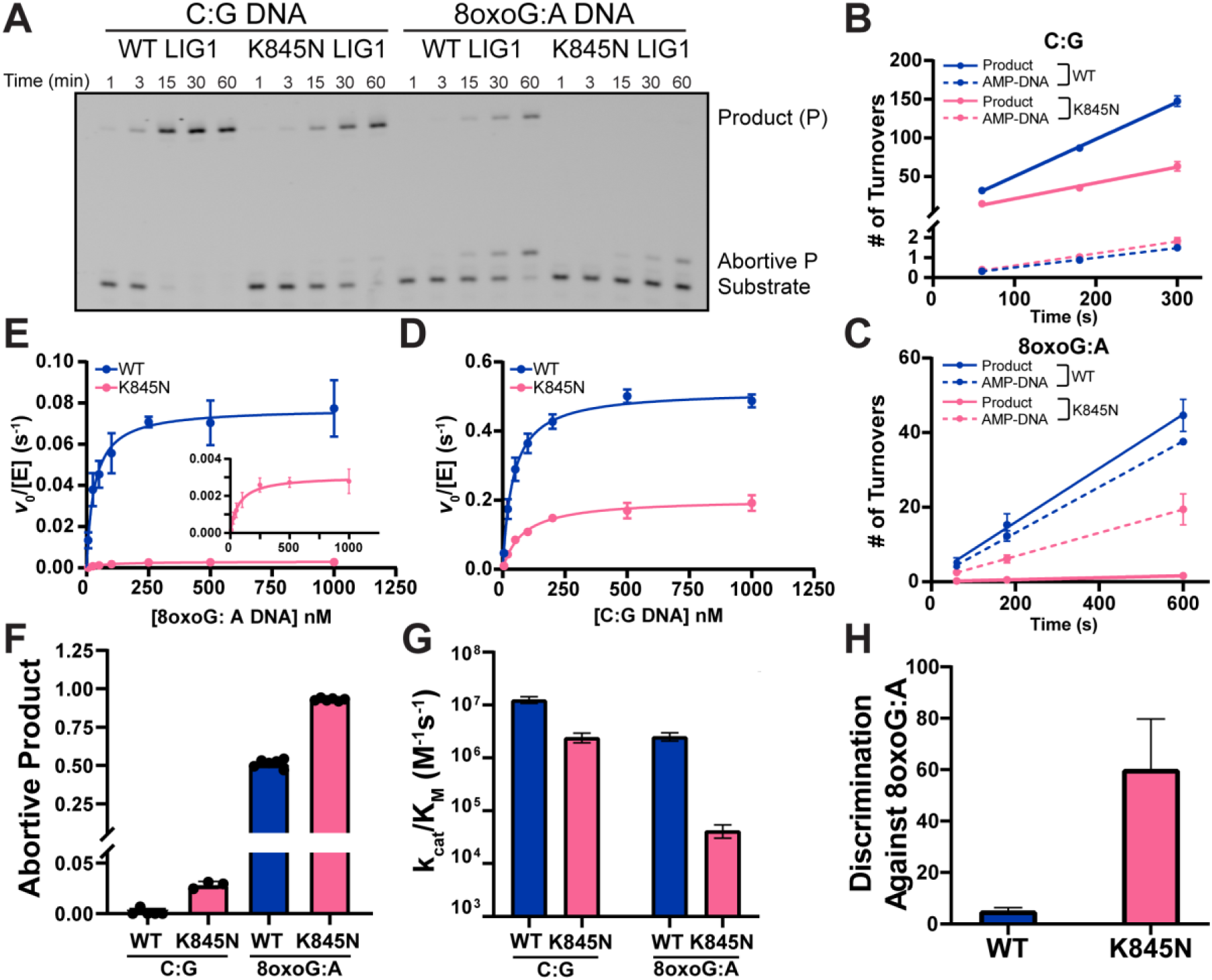
K845N is a higher fidelity, but lower activity LIG1 variant as determined by Michaelis-Menten kinetics. (A) Representative denaturing PAGE showing *in vitro* ligation reaction with WT or K845N Δ232 LIG1 (1 nM) with C:G or 8-oxoG:A containing nicked DNA (500 nM), resolving ligated product, aborted product and substrate. Reactions were performed at 37 °C with 1 mM ATP, 2 mM MgCl_2_ (1.0 mM free Mg^2^). Time courses for ligation of 1000 nM C:G (B) and 8-oxoG:A (C) 28mer substrate were plotted as number of turnovers with 1 and 5 nM LIG1 WT and 1.5 and 10 nM LIG1 K845N, respectively. Michaelis-Menten dependence for C:G (D) and 8-oxoG:A (E) 28mer nicked DNA was determined with 0.2 mM ATP and 1.0 mM free Mg^2+^. (F) The fraction abortive ligation determined from initial rates of ligation. The catalytic efficiency for ligation (G) and the discrimination against 8-oxoG:A (H) were calculated from the kinetic parameters in panels D and E. Data reported are mean ± SD (N≥3) and are summarized in *SI Appendix*, Table S2.

We next measured initial rates of reaction for the canonical nicked DNA substrate for Δ232 LIG1 WT and K845N across a range of substrate concentrations and fit the Michaelis-Menten equation to the data (Fig. 2*D*). The derived steady-state rate constants are tabulated in *SI Appendix,* Table S2. The maximal rate constant (k_cat_) values were 0.52 ± 0.01 and 0.20 ± 0.01 s^-1^ for LIG1 WT and K845N, respectively (a 2.6-fold decrease; Fig. 2*D* and *SI Appendix,* Table S2). The K_M_ value is 1.9-fold higher for K845N (*SI Appendix,* Table S2), resulting in a 4.9-fold decrease in the catalytic efficiency (k_cat_/K_M_). These k_cat_/K_M_ values are plotted in Fig. 2*G*. Analogous experiments were performed to determine steady-state kinetic parameters for the 8-oxoG:A substrate (Fig. 2*E*). There is a stark difference between the ligase activity of K845N and WT with this substrate, with most of the difference in the k_cat_ values (25-fold decrease) and a modest increase in K_M_ (2.3-fold; *SI Appendix,* Table S2), which corresponds to a 58-fold defect in k_cat_/K_M_ (Fig. 2*G*). We quantified the fraction of abortive ligation for each enzyme with each substrate (Fig. 2*F*), revealing that K845N increases the amount of abortive ligation for both substrates, with a more pronounced reduction in ligated product for the 8-oxoG:A substrate than for the undamaged substrate. The fraction of abortive ligation increases by 1.8-fold, corresponding to a 7.0-fold reduction in the fraction ligated (F_lig_) of the 8-oxoG:A substrate relative to the WT enzyme (F ^WT^/F ^K845N^ = (1 - 0.51)/(1 - 0.93) = 0.49/0.07 = 7.0). Thus, the enhanced ability of LIG1 K845N to reject the 8-oxoG:A-containing nick is attributed to both a lowered rate of nick sealing and a higher rate of abortive release of AMP-DNA.

To quantify the degree to which the K845N substitution increases the fidelity of LIG1, we calculated the discrimination factor by which Δ232 LIG1 WT and K845N distinguish 8-oxoG:A from undamaged nicks (*SI Appendix*, Table S2). Consistent with previous reports, LIG1 WT struggles to discriminate an 8-oxoG:A pair from other Watson-Crick paired nicks (25, 29, 35). This is due to the ability of 8-oxoG:A to form a Hoogsteen basepair that is further stabilized by hydrogen bonding in the LIG1 active site (25). Our data show only a 5.1-fold discrimination for LIG1 WT in ligation of 8-oxoG:A versus a canonical paired nick (Fig. 2*H* and *SI Appendix,* Table S2). In contrast, K845N discriminates against 8-oxoG:A by a factor of 60-fold, which is 12-fold higher fidelity than the LIG1 WT enzyme (Fig. 2*H* and *SI Appendix,* Table S2). K845 is highly conserved among mammalian LIG1 homologs; however, these data suggest that the K845N substitution confers unique biochemical properties. With minimal deleterious effect on ligation of canonical substrates, K845N shows an enhanced fidelity in preventing ligation of an upstream mismatch.

### Cells harboring the LIG1 K845N variant exhibit increased viability and decreased genomic DNA mutation under oxidative stress

Based on these biochemical studies, we hypothesized that the enhanced fidelity of LIG1 K845N leads to a reduction in cellular DNA mutation rate, providing a survival advantage under cellular stress conditions. Given previous data implicating oxidative stress as a potential contributor to HD (36), we tested whether K845N might protect against oxidative DNA damage-induced stress, first using HEK 293 cells stably overexpressing either full-length LIG1 WT or K845N at similar levels (*SI Appendix,* Fig. S7*A*). Cells were treated with 8 µM menadione that induces oxidative DNA damage, and viability was assessed immediately after treatment and after 24 or 48 hours of recovery (Fig. 3*A*). Immediately after treatment, viabilities (∼66%) were not different among HEK 293 cells transfected with empty vector (EV), and those expressing LIG1 WT or K845N. After 24 hours of recovery, both LIG1 WT and K845N expressing cells showed slightly increased viabilities compared to EV-transfected cells. Interestingly, after 48 hours of recovery, cells expressing K845N showed significantly higher viability than EV-transfected (*p* < 0.0001) or LIG1 WT-expressing cells (*p* < 0.0001). Furthermore, when assessed after 48 hours of recovery at various concentrations of menadione, cells expressing K845N showed consistently higher viability across all treatment conditions (*SI Appendix,* Fig. S7*B*), implying that the K845N variant confers a recovery-dependent protective effect against oxidative stress. To directly assess genomic DNA mutation rates, we performed duplex sequencing on HEK 293 cells expressing either LIG1 WT or K845N. We re-optimized conditions for a scaled-up assay that would provide sufficient cells for duplex sequencing, establishing that under these conditions, cells expressing LIG1 K845N exhibited significantly higher viability when treated with 8 µM or 20 µM menadione followed by 48 hours of recovery (Fig. 3*B*). We partitioned the cells for viability measurement and extracted DNA from the remaining cells. Duplex sequencing was performed to determine mutation frequency (MF). In both LIG1 WT- and LIG1 K845N-expressing cells MF increased in a menadione concentration-dependent manner (*SI Appendix,* Fig. S7*C*). While cells expressing LIG1 K845N exhibited a slightly higher baseline MF in the absence of treatment, they exhibited slightly lower MF in the presence of menadione. Relative to baseline (0 µM menadione), there were significant main effects (2-way ANOVA) of both menadione concentration (*p* = 0.0069) and LIG1 variant (*p* = 0.0377) on MF. Correcting for multiple comparisons revealed that the relative MF in K845N-expressing cells was significantly lower than in WT-expressing cells at 20 µM menadione (*p* = 0.0279), with a similar trend observed at 8 µM (Fig. 3*C*). These results suggest that the increased cellular viability conferred by K845N may be contributed by a decrease in mutation burden, consistent with the enhanced fidelity of K845N.

**Figure 3.**
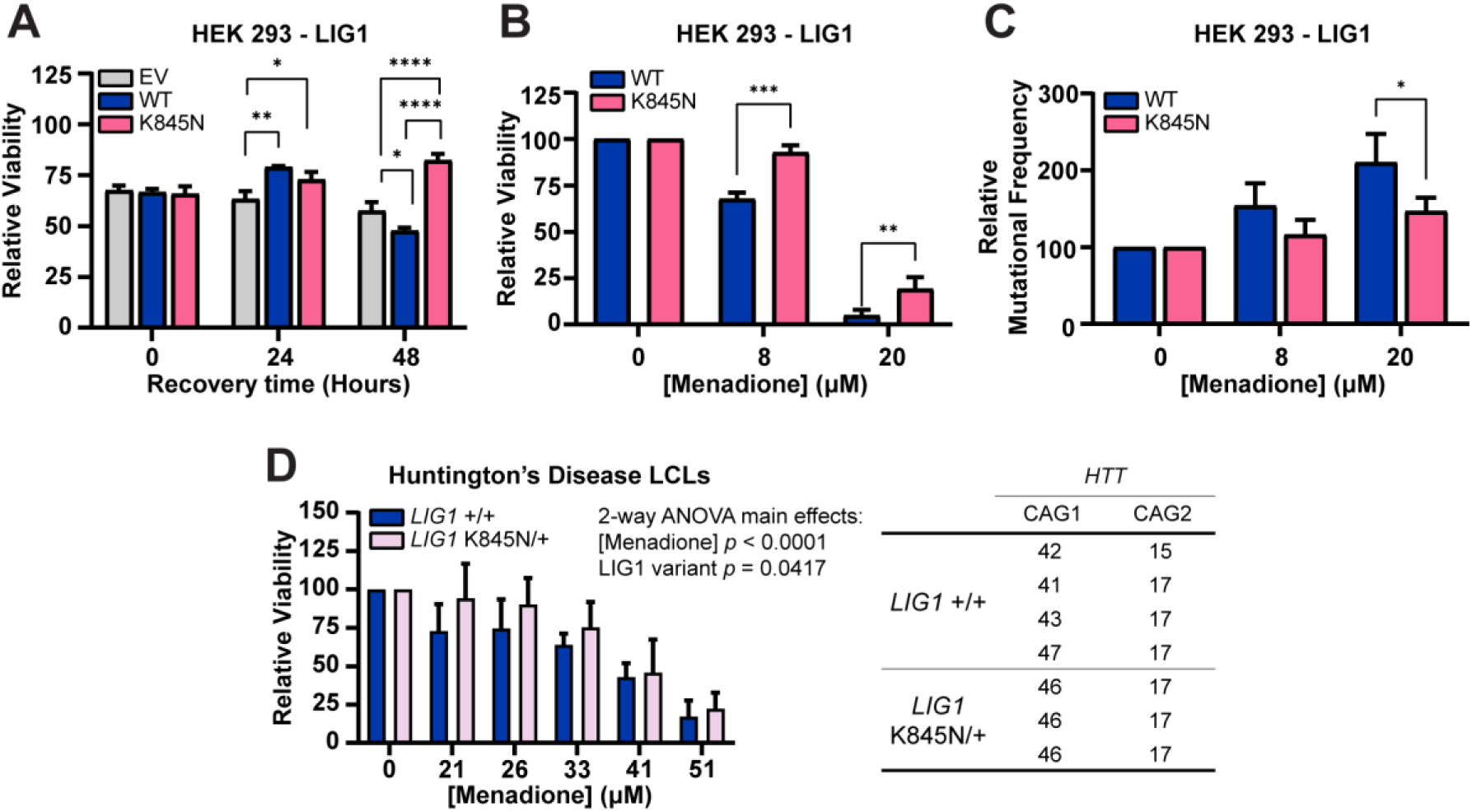
Cells harboring the LIG1 K845N variant exhibit increased viability and decreased DNA mutation under conditions that induce DNA damage. (A) HEK 293 cells (EV: empty vector-transfected or expressing LIG1 wild type (WT) or LIG1 K845N (K845N)) were treated with 8 µM menadione for 4 hours. Cells were washed in fresh medium and further incubated for the indicated times. Cellular viability was measured by CellTiter-Glo (CTG). Bar graph shows mean of 3 technical replicates ± standard deviation. Statistical significance was determined from 2-way ANOVA with Tukey’s multiple comparison test (\*\*\*\**p* < 0.0001, \*\**p* < 0.01, \**p* < 0.05). (B-C) HEK 293 cells (WT or K845N) were treated with menadione for 4 hours at the indicated concentrations and washed in fresh medium. After 48 hours recovery, the cells were harvested. (B) A portion of the harvested cells was used to measure cell viability by CTG. Bar graph shows mean % viability relative to baseline (0% menadione) of 2 technical replicates ± standard deviation (C) DNA was extracted from the remaining cells and mutation frequencies were determined by duplex sequencing. Bar graph shows mean % mutation frequency (MF) relative to baseline (0% menadione) of 2 technical replicates ± standard deviation. Statistical significance in B and C was determined from two-way ANOVA with Tukey’s multiple comparison test (\**p* < 0.05, ** *p* < 0.01, \*\*\**p* < 0.001). (D) LCLs derived from HD patients (E) were treated with menadione for 4 hours at the indicated concentrations. LCLs were either homozygous for the *LIG1* rs145821638 (GRCh38 - Chr19:48117686) reference C-allele (*LIG1+/+*) or heterozygous for the 19AM3 modifier variant A-allele (*LIG1*K845N/+). Cells were washed in fresh medium and incubated for 24 hours. Viability of each LCL was determined using CTG from the mean value of three biological replicates. Bar graph shows mean % viability relative to baseline (0% menadione) ± standard deviation (n = 4 *LIG1*+/+ independent LCLs, n = 3 independent *LIG1*K845N/+ LCLs). Statistical significance indicated in figure was determined by two-way ANOVA with Tukey’s multiple comparison test.

To extend these findings to HD patient-derived cells, we examined lymphoblastoid cell lines (LCLs) harboring only the *LIG1* rs145821638 (GRCh38 - Chr19:48117686) reference C-allele (*LIG1+/+*) or heterozygous HD LCLs harboring the 19AM3 modifier variant A-allele (*LIG1*K845N/+). LIG1 protein levels were not altered by K845N variant status or the presence of the *HTT* CAG expansion (*SI Appendix*, Fig. S8). We then treated HD patient *LIG1*+/+ (N=4) and *LIG1*K845N/+ (N=3) LCLs with varying concentrations of menadione, and as with HEK 293 cells, viability was measured immediately, and after 24 and 48 hours of recovery. While *LIG1*+/+ LCLs exhibited increased viability immediately following menadione treatment without recovery (*p* = 0.0402) (*SI Appendix*, Fig. S9*A*), *LIG1*K845N/+ LCLs showed significantly higher viability than *LIG1*+/+ LCLs after a 24-hour recovery period post-menadione treatment (Fig. 3*D*). Relative to baseline (0 µM menadione) there were significant main effects (2-way ANOVA) of both menadione concentration (*p* < 0.0001) and K845N genotype (*p* = 0.0417) on cell viability. No individual genotype comparison remained significant after multiple testing correction, though viability increased to the greatest extent in *LIG1*K845N/+ versus *LIG1*+/+ LCLs treated with 21 µM menadione (*p* = 0.0589). After 48 hours, differences were similar to those observed at 24 hours but were less distinct (*SI Appendix*, Fig. S9B). Given the small number of available HD LCLs with the K845N variant - limited to heterozygous carriers of this rare variant, and the genetic background heterogeneity of the patient samples, the statistical power in the LCLs is limited. Nevertheless, consistent with findings in HEK 293 cells, where the protective effect of LIG1 K845N emerged after a recovery period, HD LCLs expressing the endogenous LIG1 K845N variant also exhibited a recovery-dependent protective phenotype.

### The mouse K845N orthologous variant suppresses somatic CAG expansion in ***Htt*^Q111^ mice**

The somatic expansion of the *HTT* CAG repeat drives the onset of clinical phenotypes in HD. Age-dependent somatic expansion is readily measured in *Htt*^Q111^ CAG knock-in mice where it occurs at a high rate in both the striatum and the liver (18, 37). We have previously shown that somatic expansion in both these tissues can be modified to different degrees by MMR gene and various other gene knockouts and can also be modified by missense mutation impacting protein function (18, 38–41). To test whether the *LIG1* missense variant associated with delayed HD impacts somatic CAG expansion, we used CRISPR-Cas9-mediated homology-directed repair to introduce the same mutation as the human modifier (19AM3, rs145821638 C->A) into the mouse *Lig1* gene, changing the LIG1 lysine residue at amino acid 843 to an asparagine (Fig. 4*A*). As highlighted in Fig. 1*B*, K845 is a conserved residue, indicating its functional importance across species, including the mouse. We derived a *Lig1*^K843N^ line, which we crossed with *Htt*^Q111^ mice. The *Lig1*^K843N^ variant did not impact embryonic development, with *Lig1*^+/+^, *Lig1*^K843N/+^ or *Lig1*^K843N/K843N^ born at expected Mendelian ratios (*SI Appendix*, Fig. S10), contrasting with *Lig1* null alleles where homozygous null embryos do not survive beyond E15.5/E16.5 (19, 20). Consistent with the viability of *Lig1*^K843N/K843N^ mice, the K843N change did not alter *Lig1* expression levels (*SI Appendix*, Fig. S11). We also did not observe any overt phenotype in *Lig1*^K843N/+^ or *Lig1*^K843N/K843N^ mice. The mice did not develop tumors, in contrast to mice harboring the LIG1 Syndrome-associated R771W point mutation that in patient cells (4BR cell line) reduces ligase activity 20-fold (42). *Htt*^Q111/+^ mice with *Lig1*^+/+^, *Lig1*^K843N/+^ and *Lig1*^K843N/K843N^ genotypes were aged for analyses of somatic expansion. The two most unstable tissues, striatum and liver (Fig. 4*B*) revealed significantly reduced somatic expansion in *Lig1*^K843N/K843N^ striata compared to *Lig1*^+/+^ striata at 3, 6 and 10 months of age (3 months *p* < 0.001, 6 and 10 months *p* < 0.0001) and significantly reduced somatic expansion in *Lig1*^K843N/K843N^ liver at 6 and 10 months of age (6 months *p* < 0.001, 10 months *p* < 0.0001). Expansion was also significantly reduced in *Lig1*^K843N/K843N^ cortex at 3 months (*p* < 0.001) and 6 months of age (*p* < 0.0001) and in hippocampus at 6 months of age (*p* < 0.001) (*SI Appendix*, Fig. S12). There were no significant differences in the inherited repeat lengths in the cohorts of mice with different *Lig1* genotypes (*SI Appendix*, Fig. S13). Further, expansion was not impacted by the presence of a silent protospacer adjacent motif (PAM) mutation co-introduced with K843N, as determined in an independent line of mice harboring the PAM mutation in the absence of the K843N change (Materials and Methods, *SI Appendix*, Fig. S14). Of the tissues analyzed, this mutation appeared to have the most robust impact on expansion in the striatum. Although we observed *Lig1*^K843N^ allele dose-dependent trends, expansion was not significantly impacted in heterozygous *Lig1*^K843N/+^ mice relative to *Lig1*^+/+^ mice. The impact of this variant appears to be moderate, with one variant allele reducing striatal expansion by 3-5% and two mutant alleles reducing expansion by ∼15-16 % in 6 to 10-month mice. Overall, these data support reduced somatic expansion as a consequence of a LIG1 functional alteration elicited by the K843N substitution.

**Figure 4.**
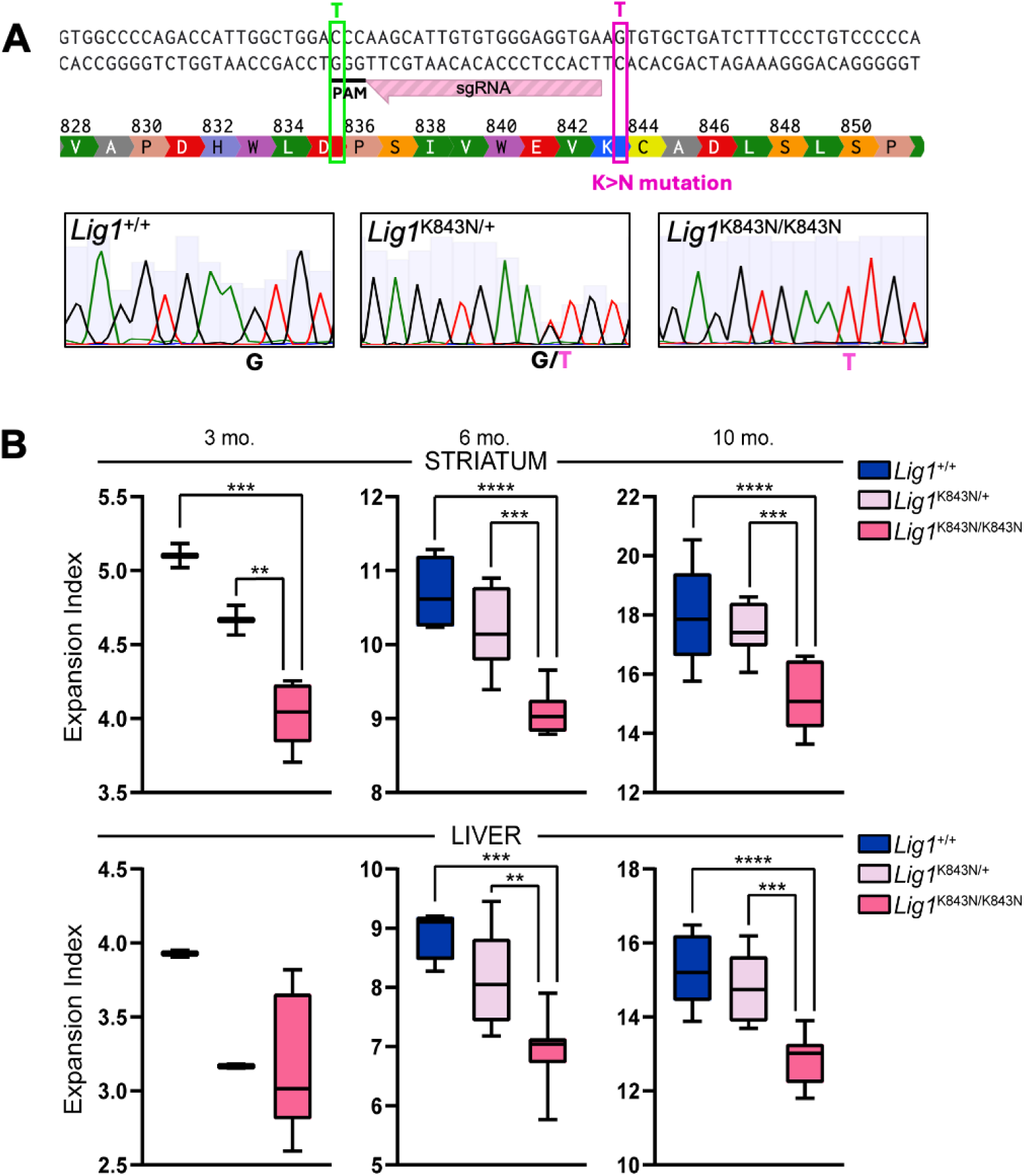
The orthologous K843N mutation suppresses somatic CAG expansion in mice. (A) CRISPR-Cas9-mediated generation of the *Lig1*^K843N^ knock-in allele. Top: The sequence is part of mouse *Lig1* exon 26 (canonical transcript ENSMUST00000177588.10) with amino acid translation showing lysine (K) at amino acid 843 (orthologous to human K845). The targeting single guide RNA (sgRNA) and protospacer adjacent motif (PAM) are shown. A 150 nt single stranded oligonucleotide (not shown – see Materials and Methods) was used for homology-directed repair to mutate G>A changing K843 to asparagine (N) (pink box). This also introduced a silent C>T change within the PAM (green box). Bottom: Sanger sequence from a *Lig1* wild-type mouse, and heterozygous and homozygous K843N knock-in mice. (B) Somatic CAG expansion indices (Min to Max box-whisker plots) from striatum and liver of *Htt*^Q111/+^ mice with different *Lig1* genotypes. 3 mo: *Lig1*^+/+^ N=2, *Lig1*^K843N/+^ N=2, *Lig1*^K843N/K843N^ N=8. 6 mo: *Lig1*^+/+^ N=4, *Lig1*^K843N/+^ N=9, *Lig1*^K843N/K843N^ N=7. 10 mo: Lig1+/+ N=9, *Lig1*^K843N/+^ N=9, *Lig1*^K843N/K843N^ N=10. \*\**p* < 0.01; *** *p* < 0.001, \*\*\*\**p* < 0.0001 (One way ANOVA, comparing all genotypes for each age and tissue, with Tukey’s multiple comparison test).

**Figure 5.**
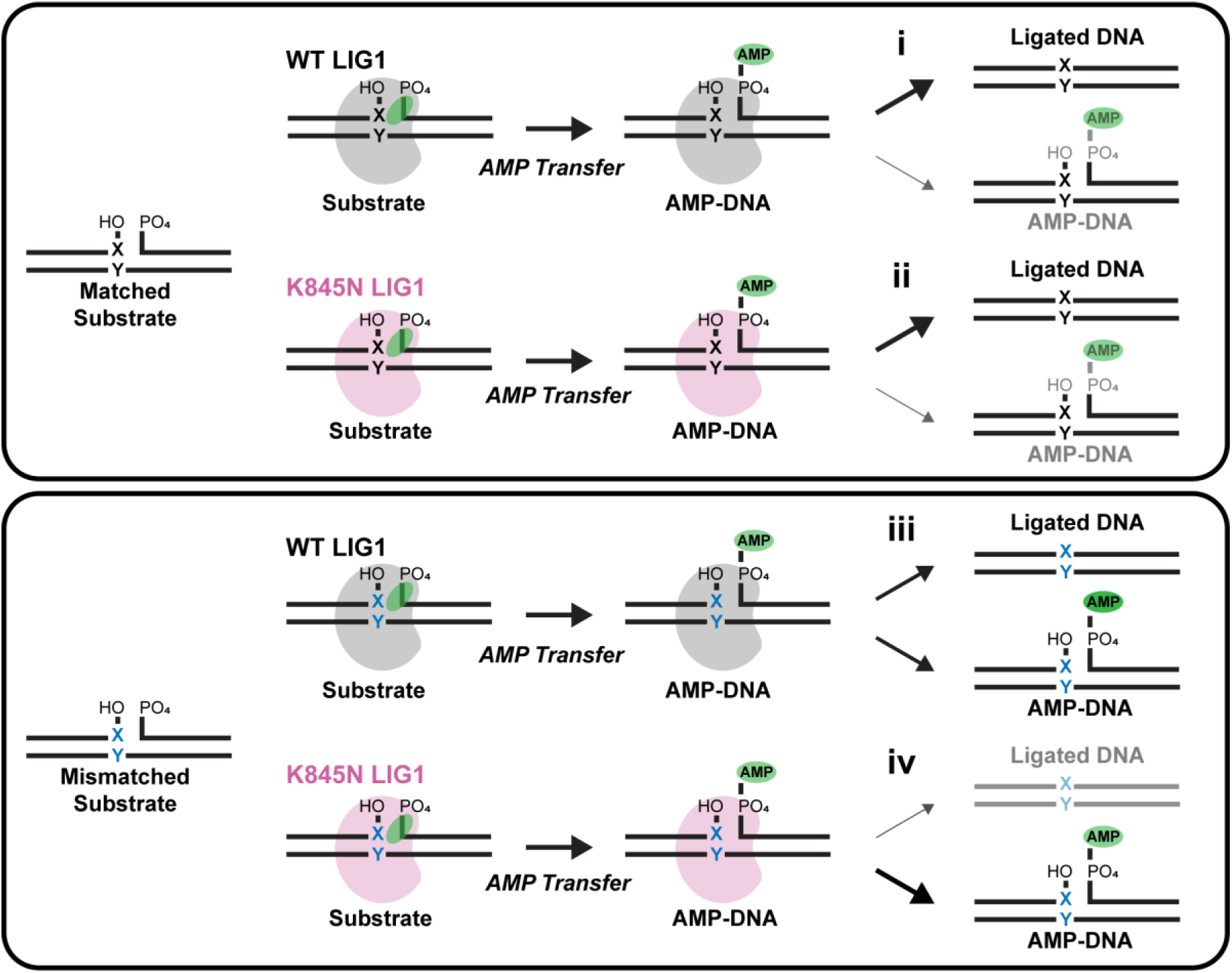
Schematic illustrating increased fidelity of ligation due to the K845N substitution. Top panel: Ligation of canonical base-paired substrate (black X:Y pair). (i) LIG1 WT produces mostly fully ligated DNA product (thick arrow and black depiction), with a small amount of aborted AMP-DNA (thin arrow and gray depiction). (ii) LIG1 K845N does not have a large impact on the proportions of ligated DNA and AMP-DNA. Bottom panel: Ligation of mismatched substrate (blue X:Y). (iii) LIG1 WT produces both fully ligated DNA product and aborted AMP-DNA (intermediate arrows). (iv) In contrast, LIG1 K845N produces less ligated product (thin arrow and gray depiction) and releases more aborted AMP-DNA (thick arrow and black depiction).

## Discussion

The minor A allele of rs145821638, coding for the *LIG1* K845N variant, was identified in GWAS as the top SNV capturing a modifier effect (19AM3) associated with delayed HD clinical phenotypes (13, 17). The impact of the modifier mechanism tagged by this SNV is substantial, delaying motor onset by 7-8 years. This rare variant, present in approximately 1 in 700 individuals (averaged across all populations described in gnomAD v.4.1.0), substitutes a conserved lysine within the OBD with a polar asparagine predicted to disrupt hydrogen bonding interactions at the domain interface between the OBD and the AdD. In this study, we evaluated the hypothesis that the HD modifier effect may reflect mechanisms impacted by this *LIG1* K845N missense change by studying its biochemical, cellular and molecular consequences.

Biallelic loss of function *LIG1* missense mutations associated with LIG1 Syndrome exhibit decreases in the maximal rate constant and in the catalytic efficiency for DNA ligation (Table 1). In comparison, our biochemical analyses of full-length and Δ232 truncated forms of LIG1 reveal the impact of K845N to be distinct from *LIG1* alleles associated with immunodeficiency syndrome (Table 1). We find that K845N maintains near-normal ligation activity on undamaged nicked DNA substrates but exhibits reduced activity toward mismatched substrates. This enhanced substrate discrimination, including against 8-oxoG:A base pairs, highlights K845N as a higher-fidelity variant of LIG1. Kinetic studies demonstrate a 59-fold reduction in catalytic efficiency (k_cat_/K_M_) on 8-oxoG:A substrates compared to WT, accompanied by an increase in abortive ligation events. Notably, the K845N variant exhibits 60-fold discrimination between mismatched and correctly paired substrates, far exceeding the 5-fold discrimination for WT.

Importantly, much of this discrimination manifests as increased abortive ligation of the improper nick which will prevent binding by another ligase and ensure additional time for other repair processes to take place. Thus, the enhanced ligation fidelity of K845N is predicted to manifest in a heterozygous background (*i.e*., even if cells express both LIG1 WT and K845N). This is relevant in HD patients where profound clinical modification is seen in individuals heterozygous for this LIG1 variant.

Functionally, this fidelity enhancement translated into a measurable benefit in cellular assays. HEK 293 cells expressing LIG1 K845N exhibit significantly increased viability in response to the oxidative damaging drug menadione relative to cells expressing LIG1 WT. Notably, this protective effect was most pronounced after an extended recovery period, suggesting that the benefits conferred by K845N may arise gradually, potentially through enhanced DNA repair processes that require time to manifest. Consistent with this, K845N-expressing cells also demonstrated a reduced mutation frequency. Supporting the relevance of this observation in a disease context, HD patient-derived LCLs heterozygous for the K845N variant showed a modest increase in viability following menadione-induced stress after recovery. Overall, our findings provide evidence that K845N can enhance genome maintenance with only modest effects on ligase activity but increased ligation fidelity.

Importantly, *Htt*^Q111^ knock-in mice carrying the orthologous *Lig1* missense variant (encoding Lig1 K843N) show a significant reduction in CAG repeat expansion in multiple tissues including the striatum and cortex that are particularly vulnerable in HD. The impact on expansion was modest; expansion was significantly suppressed in homozygous *Lig1*^K843N/K843N^ mice and trended in the same direction in heterozygous *Lig1*^K843N/+^ mice. Thus, on the face of it, the degree of expansion suppression in the mice may seem insufficient to account for the strong disease-delaying effect in patients heterozygous for this variant. However, as clinical onset is driven by somatic expansion that occurs over decades of life (8), the cumulative impact of small modifying effects may be substantial.

Indeed, a profound reduction of somatic expansion, such as that elicited by MMR gene knockouts (18), would be unexpected over a 10-month period in mice. Our study has only examined the impact of the LIG1 variant on the somatic expansion of ∼100 CAG repeats, and an interesting possibility is that its effect may depend on CAG length, for example differentially altering the recently described slow and rapid phases of somatic expansion (8). Interestingly, though *LIG1* variants modify clinical HD phenotypes, a recent GWAS of HD CAG repeat expansion in the blood did not identify significant signals at this locus suggesting that, like several other loci that impact on either clinical phenotypes or blood CAG expansion but not the other, the *LIG1* locus can involve cell-type specific effects (17). Overall, our data in mouse provide evidence that the mechanism underlying the 19AM3 disease-delaying effect may be contributed, at least in part, by an impact of the LIG1 K845N missense variant slowing the rate of somatic expansion in the brain.

It is of interest that neither knockout of *Lig1* nor the presence of the orthologous *Lig*1 R771W mutation that reduces ligase activity 20-fold had any impact on the somatic expansion of CAG/CTG repeats in mouse models of HD and myotonic dystrophy type 1 respectively, though the R771W mutation promoted maternal germline contractions (18, 42, 43). Thus, our data provide the first evidence for a role of LIG1 in the instability of CAG repeats in somatic tissues. These observations suggest that K845N has unique biochemical properties that suppress somatic CAG expansion. While the precise mechanisms of somatic repeat expansion are not fully delineated, human and mouse genetic studies as well as biochemical data support a model in which expansions are driven by the recognition and processing of extruded CAG/CTG loops/structures by the MMR machinery (17, 18, 41, 44, 45). It is currently unclear how the K845N variant of LIG1 suppresses expansion, but the elevated fidelity may mitigate expansion by rejecting substrates that support further expansion.

In summary, our data demonstrate that the K845N variant of LIG1 confers enhanced substrate discrimination and increased repair fidelity, suppresses somatic CAG expansion and is associated with increased cell survival and decreased genome DNA mutation frequency independent of repeat expansion. As somatic expansion is a key disease driver, the suppression of somatic expansion by K845N provides a plausible mechanistic underpinning for the HD-delaying 19AM3 modifier effect. Our findings also argue that the 19AM3 modifier effect may reflect, in part, increased cell survival due to genome-wide protection against DNA damage under conditions of genotoxic stress. These two potential (and non mutually-exclusive) mechanisms merit further investigation. Taken together, our results provide a mechanistic foundation for considering DNA ligase fidelity as a therapeutic target in HD and potentially in other trinucleotide repeat disorders.

## Materials and Methods

### Antibodies

The following antibodies were used: anti-LIG1 (Abcam, ab227133) and anti-alpha tubulin (Cell Signaling Technology, 3873S)

### Protein expression and purification

N-terminal FLAG-tagged full-length LIG1 WT and K845N were cloned into the pET28a vector and expressed in *Escherichia coli* BL21(DE3) at 18°C in the presence of 0.5 mM Isopropyl β-D-1-thiogalactopyranoside (IPTG) for 18 hours. Cells were harvested in 500 mM NaCl, 50 mM Tris pH 8.0, 1 mM EDTA, 5 % Glycerol in presence of cOmplete™, Mini, EDTA-free Protease Inhibitor Cocktail (Roche). The cells were lysed, and the lysates were cleared by centrifugation at 15,000 rpm for 40 minutes. The supernatant was incubated with M2-affinity resins (Sigma-Aldrich) for 4 hours and the proteins were eluted with 0.4 mg/mL FLAG peptide. The eluted samples were dialyzed into 150 mM NaCl, 50 mM Tris pH 8.0, 5% Glycerol and concentrated using Amicon Ultra centrifugal filter devices (Millipore).

Δ232 LIG1 was expressed from a pET19 plasmid vector (24), and the K845N variant was created using site-directed mutagenesis. Sequences were confirmed by full plasmid sequencing. LIG1 variants were expressed via auto-induction in TB + 0.2x trace metals, 0.2% lactose, 0.05% glucose, 0.4% glycerol and 5 µg/mL carbenicillin (46). After 4 hours of shaking at 37 °C, cells were pelleted and diluted with an equal volume of lysis buffer (50 mM Tris-Cl pH 7.5, 10% glycerol, 300 mM NaCl, 5 mM β-mercaptoethanol) and protease inhibitors were added (0.5 mM PMSF, 0.5 µg/mL leupeptin, 0.7 µg/mL pepstatin A, 0.01% IGEPAL). Cells were then lysed with a cell homogenizer, and collected via centrifugation (19,000 rpm, 30 minutes at 4 °C with an SS-34 rotor). PEI precipitation removed excess nucleic acids (0.1% poly(ethyleneimine) *v/v* cell supernatant while stirring at 4 °C). After the soluble fraction was centrifuged again, LIG1 was purified via standard low-pressure His-Trap column. Briefly, a 5 mL Ni-NTA column was first washed with a high-salt buffer (20 mM HEPES pH 7.5, 20 mM imidazole, 500 mM NaCl), followed by a low-salt buffer (20 mM HEPES pH 7.5, 50 mM imidazole, 100 mM NaCl). The soluble fraction was then loaded onto the column, followed by a low-salt buffer wash. Finally, proteins were eluted into fractions via a gradient between low-and high-imidazole buffers (20 mM HEPES pH 7.5, 50-300 mM imidazole, 100 mM NaCl). LIG1 adenylation and His-tag cleavage took place overnight at 4 °C with 1 mM ATP, 11 mM MgCl_2_ 0.5 mM EDTA and purified precision protease enzyme. Next, LIG1 was further purified via FPLC HiTrap Q column using low salt (20 mM HEPES pH 7.5, 1 mM EDTA, 5 mM β-mercaptoethanol) wash buffer and high salt (20 mM HEPES pH 7.5, 1 mM EDTA, 2 M NaCl, 5 mM β-mercaptoethanol) elution buffer using a buffer gradient. LIG1 was then purified again via HiTrap Blue column using the same buffers and method as the HiTrap Q column. Finally, purified Δ232 LIG1 variants were buffer exchanged into storage buffer (25 mM Tris-Cl pH 7.5, 150 mM NaCl, 0.1 mM EDTA, 1 mM DTT) and purity was confirmed via SDS-PAGE.

### Preparation of DNA substrates

The DNA oligonucleotides were synthesized by Integrated DNA Technologies (IDT) and gel purified. Concentrations were determined by the absorbance using the predicted A_260_ values. The nicked DNA substrates were generated by annealing the oligos in a 1:1.5:2 ratio (10 µM PO4 to 15 µM template to 20 µM OH) in buffer comprised of 10 mM NaMES (pH 6.5) and 50 mM NaCl. The mixture was incubated at 95 °C for 5 minutes and then gradually cooling the solution to 4 °C, decreasing the temperature by 1°C every 5 seconds. Annealed substrates were stored at 4°C. The sequences of oligonucleotide substrates are listed in *SI Appendix*, Table S1.

### Active site titration

To determine the active concentration of LIG1 in solution, active site titration assays were utilized (34). Briefly, full-length (12.5 nM to 400 nM) and Δ232 (20 nM to 300 nM) LIG1 variants were combined with 100 nM annealed assay substrate in the absence of ATP to prevent multiple turnover. This reaction was conducted in standard reaction buffer containing 50 mM MOPS pH 7.5, 1 mM DTT, 0.5 mg/mL BSA, 10 mM MgCl_2_, with the ionic strength adjusted to 150 mM using NaCl. After incubation at 37 °C for 1 hour, reactions were quenched with standard loading buffer (90% formamide, 50 mM EDTA, 0.01% bromophenol blue, 0.01% xylene cyanol) and were heated to 95 °C for 5 minutes. The sealed product was separated from the substrate by denaturing PAGE on a 15% polyacrylamide/8 M urea gel. Active enzyme concentration was determined by segmental linear regression analysis in GraphPad Prism software.

### Gel-based ligation assays for full-length LIG1

The ligation assays for full-length LIG1 were performed in 150 mM NaCl, 50 mM MOPS (pH 7.5), 1.0 mM DTT, 0.5 mM ATP, 10 mM MgCl_2_, 300 nM nicked DNA substrate. Purified LIG1 (final 5 – 40 nM) was added to ligation buffer and the mixture was incubated for 5 minutes at 37 °C. Ligation reactions were quenched by cooling to 4°C followed by addition of loading buffer (90 % formamide, 50 mM EDTA), heated at 95°C for 5 minutes. The samples were separated on 15 % denaturing urea acrylamide mini gels and the gel was scanned with iBright imaging system (Thermo Fisher Scientific). Fluorescence intensity was analyzed using image Lab software (Bio-Rad). The fraction of observed ligation product was determined by dividing band intensity of product by the total band intensity.

### Steady-state kinetics with Δ232 LIG1

Steady-state kinetics experiments were performed with Δ232 LIG1 at 37 ^°^C in standard reaction buffer with 1.2 mM MgCl_2_ (1 mM free Mg^2+^), 0.2 mM ATP, 50 mM MOPS pH 7.5, ionic strength adjusted to 150 mM using NaCl, and varying LIG1 and DNA concentrations. Each reaction time point was quenched in standard loading buffer and heated to 95 °C before gel loading. Sealed and abortive products were separated from substrate by denaturing PAGE on 15% polyacrylamide/8 M urea sequencing gels. Fluorescent bands were detected using an Amersham Typhoon 5 imager (GE) and quantified with ImageQuant TL software (GE).

Initial rates were determined by fitting the fraction product (< 0.20) to straight lines in GraphPad Prism, calculating reaction velocity from the slope. The fraction of abortive ligation is defined by Eq. 1. The initial rates of sealed product formation were fit with the Michaelis-Menten equation (Eq. 2) to determine *k*_cat_ and K_M_ values, as well as the catalytic efficiency (*k*_cat_/K_M_). The discrimination between 8-oxoG:A-containing substrate and undamaged (C:G) substrate is given by Eq. 3. In some cases, the time courses are expressed as the number of turnovers, which is calculated by dividing the concentration of ligated product by the concentration of enzyme.

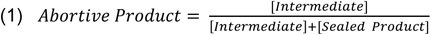

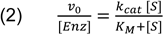

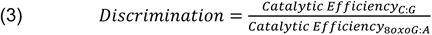

### HD patient and control LCLs

We used a panel of LCLs from seven previously collected and genotyped HD individuals participating in HD research. Of these, three were heterozygous for the 19AM3 modifier variant rs145821638 (GRCh38 - Chr19:48117686-C-A), all with *HTT* CAG repeat lengths 46/17, and four lacked the minor (A) allele, with *HTT* CAG repeat lengths 42/15, 41/17, 43/17 and 47/17. We also analyzed LCLs from two unaffected individuals lacking the 19AM3 minor allele (*HTT* CAG repeat lengths 17/15 and 20/19). LCLs were obtained from the MGB IRB-approved CHGR Neurodegenerative Repository. Genotyping of the normal and expanded CAG repeats was performed using a standardized PCR assay (47). *LIG1* SNV rs145821638 (GRCh38 - Chr19:48117686-C-A) genotype was determined using a TaqMan SNP genotyping assay from Life Technologies. The reaction contained 1x TaqMan Genotyping Master mix (Life Technologies, part # 4371355) 1x of custom TaqMan SNP Genotyping assay mix (Assay mix comes at 40X is diluted to 20X in TE buffer pH8.0 (Life Technologies part #AM9849)) and 10 ng of DNA in total reaction volume of 12.5 ul. The reaction was run on the Roche LightCycler 480 II with the following conditions: 95°C for 10 mins, followed by 35 cycles of 92°C for 15 seconds, 60°C for 1 minute. Data was collected at the end of each amplification cycle and known homozygous wildtype and heterozygous controls were run with every plate to assign alleles.

### Cell viability assay and duplex sequencing

To generate HEK 293 cells stably overexpressing either full-length LIG1 WT or K845N, HEK 293 cells were transfected with pcDNA3.1-N-terminal FLAG-tagged full-length LIG1 WT, pcDNA3.1-N-terminal FLAG-tagged full-length LIG1 K845N or pcDNA3.1-Empty vector in a 24 well plate with Lipofectamine 3000 (Life Technologies). The K845N construct was generated by site-directed mutagenesis kit (New England Biolabs) of the WT pcDNA3.1-LIG1 plasmid. Following transfection, cells were selected with G418 (Geneticin) and subsequently sorted to establish stable cell lines. They were seeded into 96 well plates at a density of 2 x 10^4^ cells/well in DMEM medium, and incubated overnight at 37°C. The cells were treated with multiple dilution of menadione (MP biomedicals) for 4 hours, washed in a fresh medium and incubated for indicated durations at 37°C with 5 % CO_2_. The number of viable cells was measured using the CellTiter-Glo (CTG) cell viability assay (Promega) according to the manufacturer’s instructions. The data were expressed as the percentage of the amount of ATP measured relative to untreated controls. For duplex sequencing, HEK 293 cells were seeded into 6 well plates at a density of 8 x 10^5^ cells/well in DMEM medium, and incubated overnight at 37 °C. The cells were treated with 8 and 20 μM of menadione (MP biomedicals) for 4 hours, washed in a fresh medium and incubated for 48 hours under normal conditions. After incubation, cells were harvested. A portion of the harvested cells was taken and cell viability was measured using CTG. The remaining cells were pelleted by centrifuge and stored at -80°C until analysis. The cell pellets were sent to inotiv for DNA extraction, library preparation, duplex sequencing and analysis of mutation frequency. Briefly, the TwinStrand Duplex Sequencing™ Mutagenesis Panel (Human-50), v2.0 kit (TwinStrand Bioscience, Seattle, WA, USA) was used for DNA library preparation. All libraries were pooled together at equimolar concentrations and loaded on 1 lane of a 25B (2 × 150 bp) flow cell, to target ∼542 million paired-end reads per sample (assuming capacity of the flow cell is 52 billion paired-end reads), which is expected to yield ∼1–1.25 billion duplex bp per sample. All libraries were sequenced using the Illumina NovaSeq X Plus platform.

LCLs were grown in humidified incubator in RPMI 1640 medium (Sigma-Aldrich) supplemented with 10% fetal bovine serum and 1% Penicillin-Streptomycin-Glutamine at 37°C and 5% CO_2_. LCLs were seeded into 24 well plates at a density of 5 x 10^5^cells/well in RPMI 1640 medium (Gibco) and treated with multiple dilution of menadione (The highest concentration was 100 μM and serial dilution of 4/5 were made) for 4 hours, after which the medium was replaced with fresh medium for indicated recovery periods. The number of viable cells was determined using CTG (Promega) in accordance with the manufacturer’s instructions. The viability was expressed as the percentage of the amount of ATP measured relative to untreated controls.

### Generation of *Lig1*^K843N^ knock-in mice

This study was carried out in accordance with the recommendations in the Guide for the Care and Use of Laboratory Animals of the National Institutes of Health under an approved protocol of the Massachusetts General Hospital Institutional Animal Care and Use Committees (IACUCs) (MGH: protocol 2009N000216). All animal procedures were carried out to minimize pain and discomfort, under the approved IACUC protocol. Animal husbandry was performed under controlled temperature, humidity and light/dark cycles. CRISPR/Cas9-mediated homology-directed repair was carried out essentially as reported (48) (Genome Modification Facility of Harvard University). Candidate single guide RNAs (sgRNAs) were chosen using the Broad Institute design tool (49, 50), four sgRNAs screened in embryos using Sanger sequencing to test guide efficiency, from which one was picked based on having the highest estimated editing efficiency. This gRNA (5’>3’ 5’ TCACCTCCCACACAATGCTT; GRCm39 chr7:13,043,112-13,043,131 [-strand]) was co-injected into C57BL/6J zygotes with Cas9 protein (Alt-R® S.p. Cas9 Nuclease V3, IDT) and the following 150 nt single stranded oligonucleotide donor (ssODN) template (Genewiz) complementary to the guide sequence.

5’ TCTGCTCACTGCAGGCCCTGGTATTGCCTACCCCACGCCCCTATGTGAGGATTGAT GGGGCAGTGGCCCCAGACCATTGGCTGGA**T**CCAAGCATTGTGTGGGAGGTGAA**T**T GTGCTGATCTTTCCCT The ssODN harbored the coding G>T mutation (red font) specifying asparagine at position 843 in the mouse LIG1 protein (in *Lig1* exon 26, canonical transcript ENSMUST00000177588.10) and a silent G>T mutation (blue font) within the S.p. Cas9 PAM sequence (converting the GGG PAM on the minus strand to GGA) to prevent endonuclease cleavage of the recombined sequence. Injected zygotes were implanted into pseudopregnant females and pups screened at weaning by Sanger sequencing for the presence of the targeted allele. We established germline transmission from a pup that exhibited ∼100% of targeted sequence and further verified the presence of the targeted mutations (K843N and silent PAM) and absence of other sequence variants using MiSeq (primers F: 5’-CAGGCCCTGGTATTGCCTAC; R: 5’-CCATCACCTCTGCCTTCCTT). The line was subsequently maintained in crosses with C57BL/6J and sperm frozen at the Jackson laboratories. We also serendipitously obtained a line harboring only the silent PAM mutation, which we maintained in the same way and used as an additional control line. Mice were genotyped using custom designed TaqMan SNP genotyping assays from Life Technologies, specific for either the K843N variant or the PAM sequence variant. Reactions contained 1x TaqMan Genotyping Master mix (Life Technologies, part # 4371355) 1x of custom TaqMan SNP Genotyping assay mix (Assay mix comes at 40X is diluted to 20X in TE buffer pH 8.0 (Life Technologies part #AM9849) and 10 ng of DNA in total reaction volume of 12.5 µl. The reaction was run on the Roche LightCycler 480 II with the following conditions: 95°C for 10 min, followed by 35 cycles of 92°C for 15 seconds, 60°C for 1 minute, and data collected at the end of each amplification cycle. Known wild-type, heterozygous and homozygous mutant controls were run with every plate to assign alleles.

### Mouse crosses with *Htt*^Q111^ and measurement of somatic CAG expansion

*Htt*^Q111^ mice on the C57BL/6J genetic background (51) were crossed with *Lig1^K843N^* mice to generate *Htt*^Q111/+^ mice with *Lig1*^+/+^, *Lig1*^K843N/+^ and *Lig1*^K843N/K843N^ genotypes. Separate crosses were set up between *Htt*^Q111/+^ mice and the “PAM only” mutant line to generate a small cohort of *Htt*^Q111/+^ PAM/PAM control mice. *Lig1* alleles were genotyped as above and the *Htt*^Q111^ allele genotyped according to Kovalenko et al (52). Genomic DNA from mice at 3, 6 or 10 months of age was isolated from tissues using the DNeasy DNA blood and tissue kit (Qiagen) and somatic CAG instability analysis was performed using a human-specific PCR assay that amplifies the *HTT* CAG repeat from the knock-in allele (40). The forward primer was fluorescently labeled with 6-FAM (Thermo Fisher Scientific) and products were resolved using the ABI 3730xl DNA analyzer (Thermo Fisher Scientific) with GeneScan 500 LIZ as internal size standard (Thermo Fisher Scientific). GeneMapper v5 (Thermo Fisher Scientific) was used to generate CAG repeat size distribution traces. CAG expansion indices were calculated from GeneMapper peak height data as previously described, using a 5% relative peak height threshold cut-off (53). Expansion indices in all tissues were determined relative to the modal allele in the liver trace from the respective mouse (representing the modal repeat length in a population of liver cells that is very stable over time and distinguished as mice age from the unstable hepatocyte population (37).

### Assaying *Lig1* expression in mice

Total RNA was extracted from 25-30 mg of mouse liver using a hybrid Trizol-Qiagen method (modified from https://imageslab.fiu.edu/wp-content/uploads/2020/02/RNA-extraction_Trizol_RNeasy_hybrid.pdf). The tissue was homogenized in Trizol reagent followed by phase separation according to Trizol manufacturer’s protocol. The top (aqueous) phase was mixed with 2 parts of RLT Plus buffer and applied to genomic DNA Eliminator spin columns (both buffer and columns were from the RNeasy Plus Mini kit, Qiagen, 74134). All the following steps were performed according to the RNeasy Plus Mini kit protocol. Quality control of resulting RNA was done with Agilent TapeStation 3000. The cDNA was synthesized with the Superscript III kit (Invitrogen, 18080-051) using oligo(dT) primer, according to the manufacturer’s instructions. The relative expression level of *Lig1* was quantified by PCR, using 2 µl of undiluted cDNA, 18 µl Taqman FastAdvanced Master Mix (Invitrogen, 4444556), and 1 µl of a ready-to-use Taqman assay from Invitrogen (*Lig1*: Mm00495331_m1 mapping to exons 21-22 and Mm01199942_m1 mapping to exons 3-4). PCR conditions were as follows: UNG incubation: 50°C (2 min), initial denaturation and polymerase activation: 95°C (20 s), 40 cycles of 95°C (3 s), 60°C (30 s), final extension 72°C (10 min). Relative expression values (DC_t_) of *Lig1* for each mouse were calculated using geometric mean of the C_t_ values of three housekeeping genes (*Ppia*, Mm02342430_g1; *Actb*, Mm00607939_s1; *Gusb*, Mm00446953_m1). Expression fold change (2^-DDCt^) of *Lig1* for each mouse was calculated relative to the geomean of relative expression values (DC_t_) of the control group (*Lig1^+/+^*) mice.

## Statistical analyses

Statistical analyses were performed using GraphPad Prism v10. For comparisons of menadione-induced effects between LIG1 WT and LIG1 K845N-expressing cells, two-way ANOVA followed by Tukey’s multiple comparison test were conducted. This analysis was applied to both cell viability assays and mutation frequency measurements obtained from duplex sequencing. Comparisons of somatic expansion and *Lig1* expression levels in mouse tissues were carried out using one-way ANOVA with Tukey’s multiple comparison test. Mendelian ratios of pups with different *Lig1* genotypes were assessed with Chi-squared tests. Significance was represented using a symbol-based threshold system: \*\*\*\**p* < 0.0001, \*\**p* < 0.001, \*\**p* < 0.01, and \**p* < 0.05

## Acknowledgments

This work was supported by NIH grant R01NS127866 (V.C.W and I.S.S), R01 NS049206 (V.C.W), R01 NS114065 (I.S.S.), R35GM149546 (P.J.O.), NS091161 (J.F.G.) and the CHDI Foundation (V.C.W, I.S.S, M.E.M, J.F.G). D.H.B was supported in part by an NSF GRP (DGE-1841052). We are grateful to Dr. Lin Wu of the Harvard Genome Modification Facility for her input.

## Author Contributions

E.L., W.K., D.H.B., Y.L., M.K., F.S., E.O., R.M., M.A.A., T.G., B.D, B.S, J.R and R.M.P. designed and/or conducted experiments and/or performed data analysis. S.K, R.L, D.L, M.E.M and J.F.G provided resources. P.J.O., V.C.W and I.S.S. contributed to the design of the study. E.L., D.H.B., P.J.O., V.C.W. and I.S.S. were involved in writing the manuscript, which was reviewed by all the authors.

