## Supplemental Information_Figures and Tables for "Huntington’s disease LIG1 modifier variant increases ligase fidelity and suppresses somatic CAG repeat expansion"

### Supplementary Figures

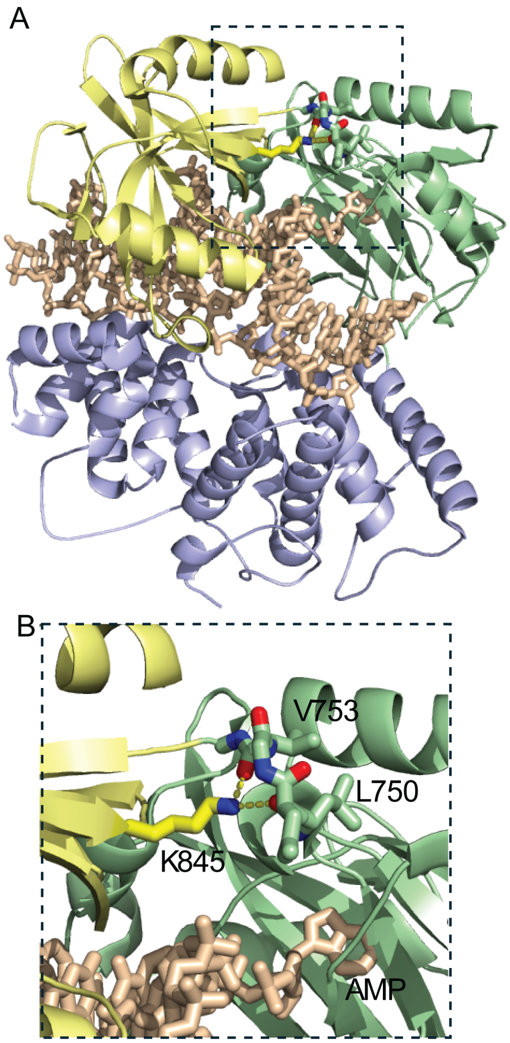


**Figure S1. Interdomain contacts made by K845 from the OBD (yellow) to the AdD (green).**

(A) LIG1 WT in complex with adenylylated DNA rendered from PDB 6p0c [25]. With the DBD (blue), the three domains of LIG1 encircle the nicked DNA. (B) Close-up view of K845 interactions. The epsilon amino group of K845 makes bidentate hydrogen bonds with the backbone amides of L750 and V753 (2.4 and 2.7 Å respectively, yellow dashed lines). K845N forms a lynchpin at the interface between the OBD and AdD.

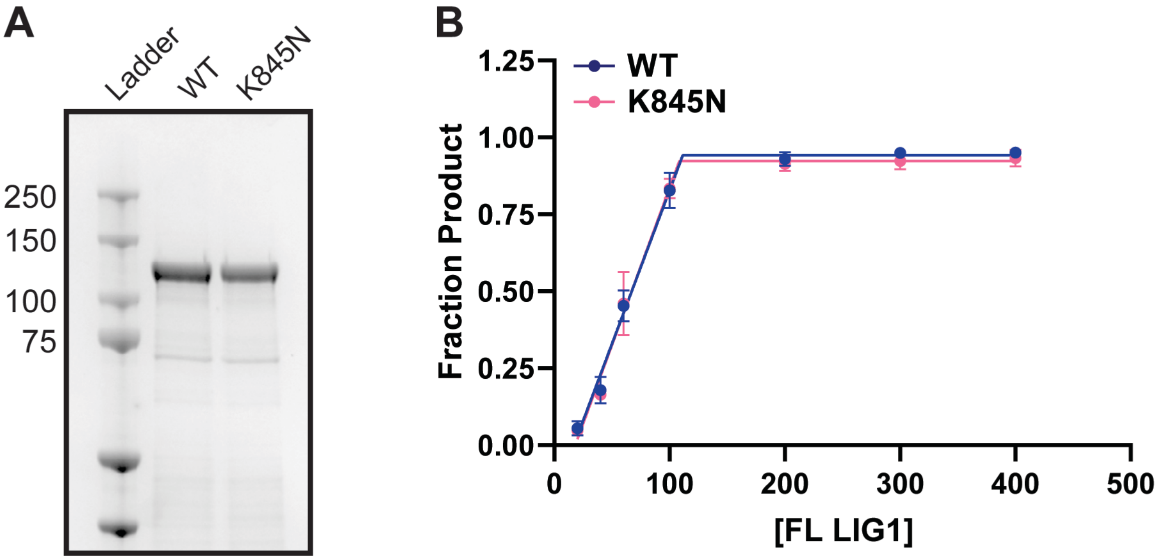


**Figure S2. Purification and active concentration of full-length LIG1 WT and K845N.**

(A) Full-length LIG1 WT and K845N proteins were analyzed by 4-12 % SDS-PAGE stained with Coomassie blue. The impurities seen in the gel appear to be at similar levels in WT and K845N preparations. (B) Active site titration assays were performed to measure the concentration of active LIG1 using 100 nM of A:T (34mer) nicked DNA substrate in the absence of ATP. Reactants were analyzed in 15 % TBE-Urea polyacrylamide gels (not shown). The line graphs show the quantification of the fraction of ligated product from three independent experiments (mean ± SD). WT and K845N were determined to be 89 ± 6% and 92 ± 6% active, respectively.

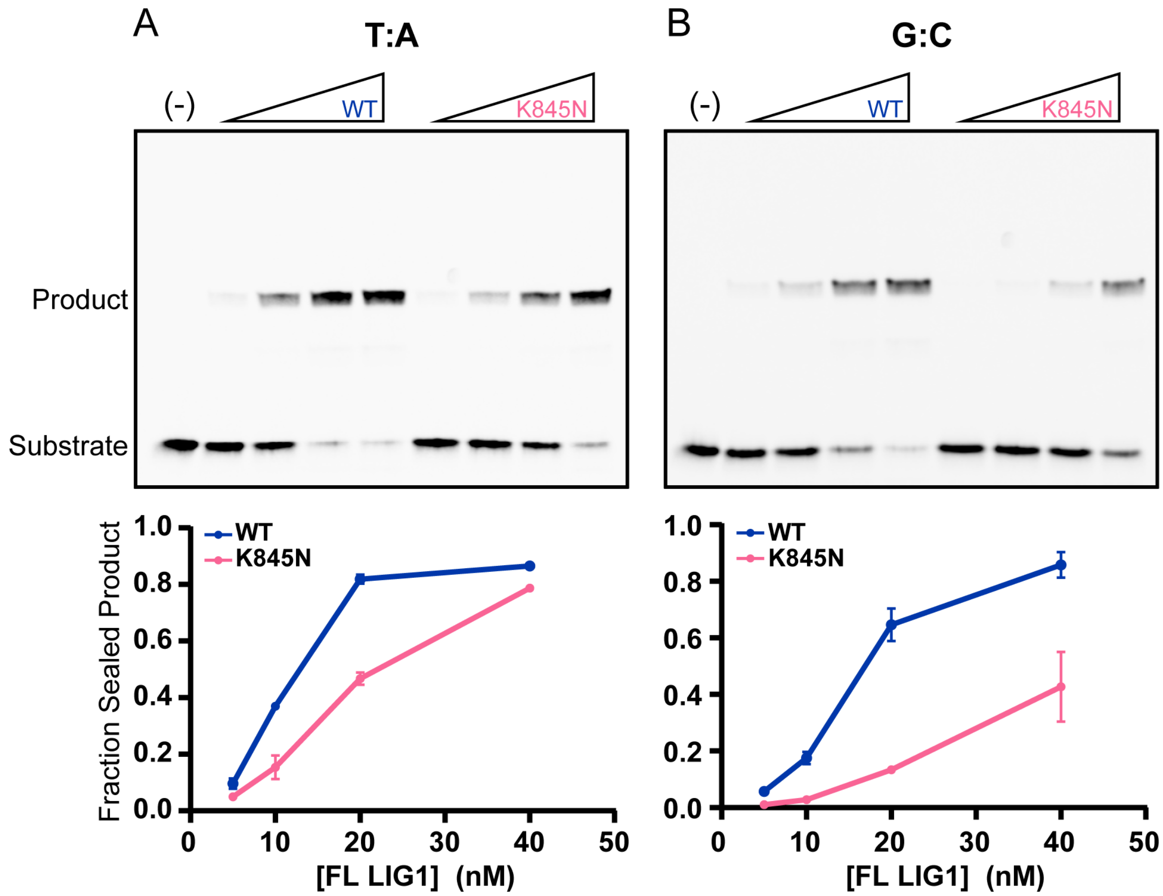


**Figure S3. Ligation of canonical substrate by LIG1 WT and K845N**

Representative gels showing the ligase activity of LIG1 WT and K845N. T:A (A) or G:C (B) containing nicked DNA (34mer; 300 nM) was incubated with increasing amounts of the proteins for 5 min at 37 °C. Reactants were analyzed in 15 % TBE-Urea polyacrylamide gels. Reactions were performed with 1 mM ATP, 10 mM $MgCl_2$, 50 mM MOPS pH 7.5 and 150 mM NaCl. The line graphs show the quantification of the fraction of ligated product from three independent experiments (mean ± SD). Note that in panel (A), the error bars for the WT group are smaller than the symbol size and may not be visible.

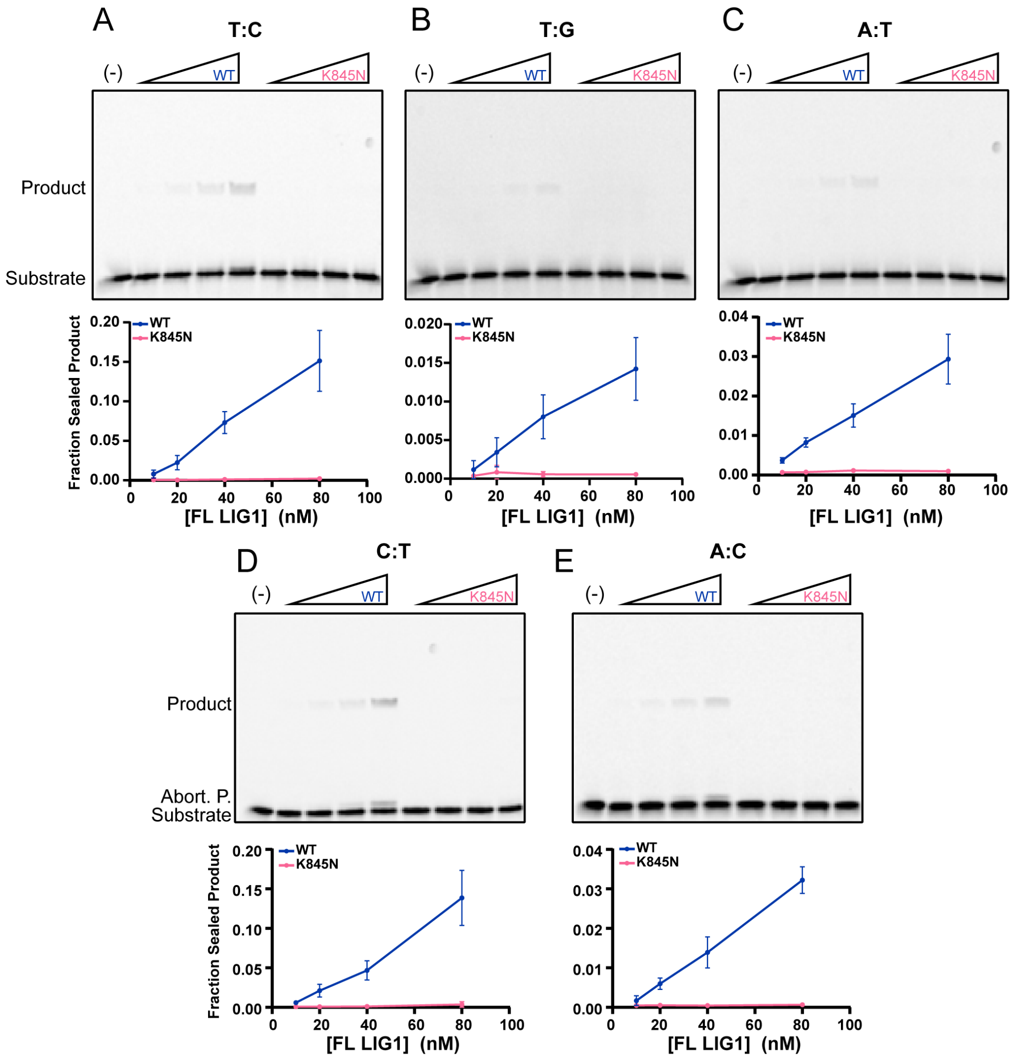


**Figure S4. Ligation of mismatched substrate by LIG1 WT and K845N**

Representative gels showing the ligation of mismatched DNA substrates by LIG1 WT and K845N. T:C (A), T:G (B), C:A (C), C:T (D), and A:C (E) containing nicked DNA (34mer) was incubated with increasing amounts of enzyme for 5 min at 37 °C. Reactions were performed with 1 mM ATP, 10 mM $MgCl_2$, 50 mM MOPS pH 7.5 and 150 mM NaCl. Reactants were analyzed by 15 % TBE-Urea polyacrylamide gels. The line graphs show the quantification of the fraction of ligated product from three independent experiments (mean ± SD).

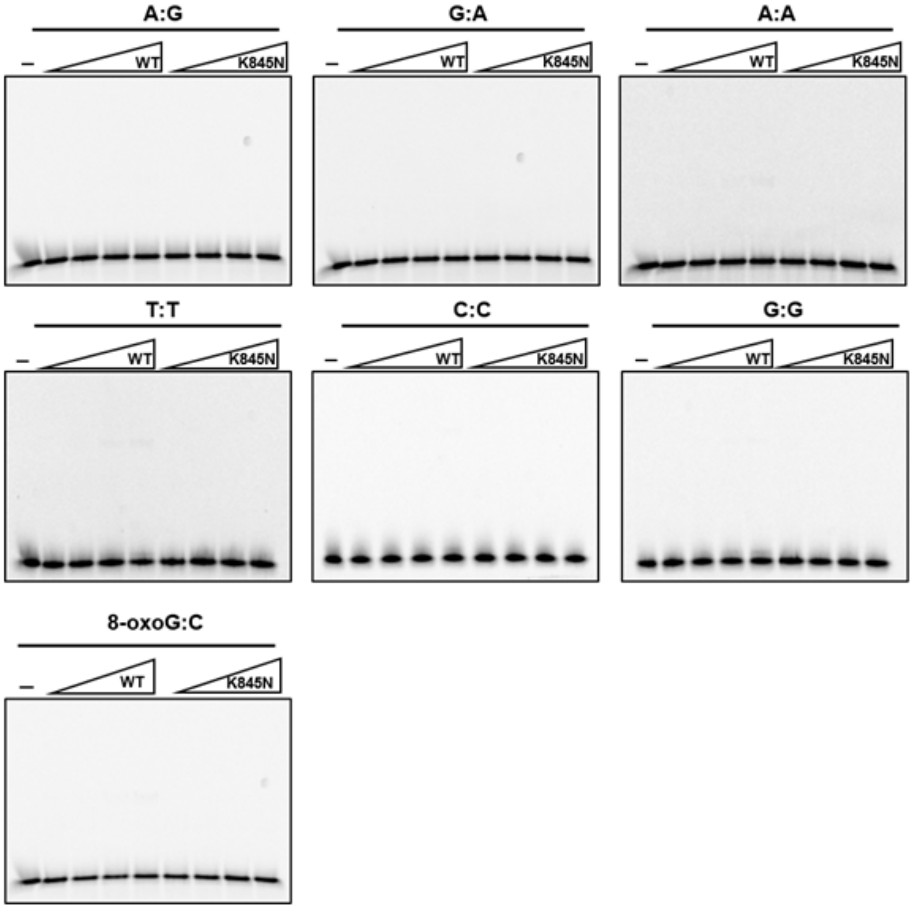


**Figure S5. Ligase assays with additional mismatched substrates.** Representative gels showing the absence of significant ligated product for several different 3' X:Y 34mer contexts. DNA (34mer; 300 nM) was incubated with 10-80 nM enzyme for 5 min at 37 °C. There was no detectable product formed by either LIG1 WT or K845N, and therefore no quantitation was performed.

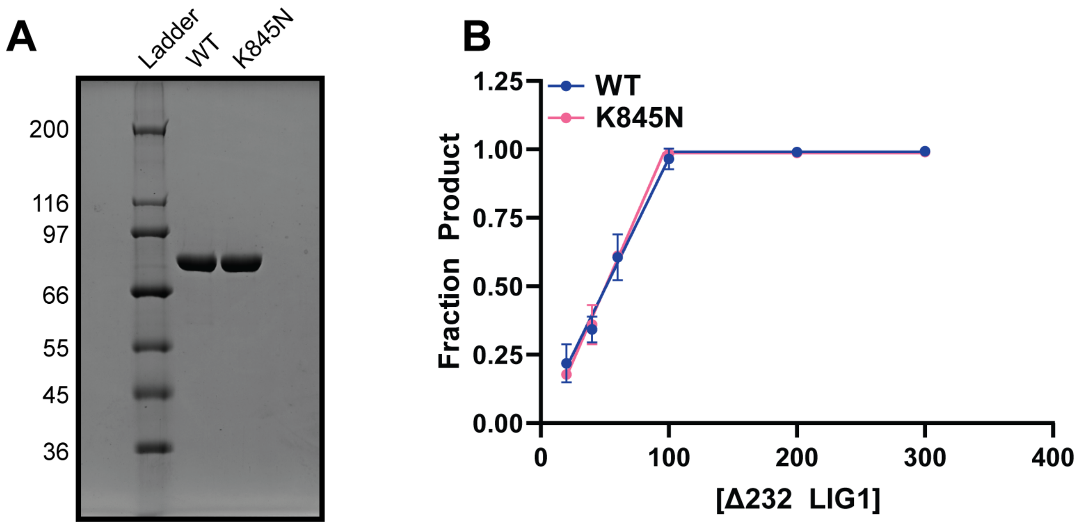


**Figure S6. Purification and active concentration of WT and K845N ∆232 LIG1.**

(A) Purified LIG1 WT and K845N proteins (1 µg per lane) were analyzed with 10 % SDS-PAGE (Δ232 LIG1 is 76.5 kDa). (B) An active site titration assay was conducted in the absence of ATP with 100 nM nicked DNA substrate (C:G 28mer) to determine the concentration of active LIG1 in solution. LIG1 WT and K845N were determined to be 97 ± 8 % and 104 ± 7 % active, respectively.

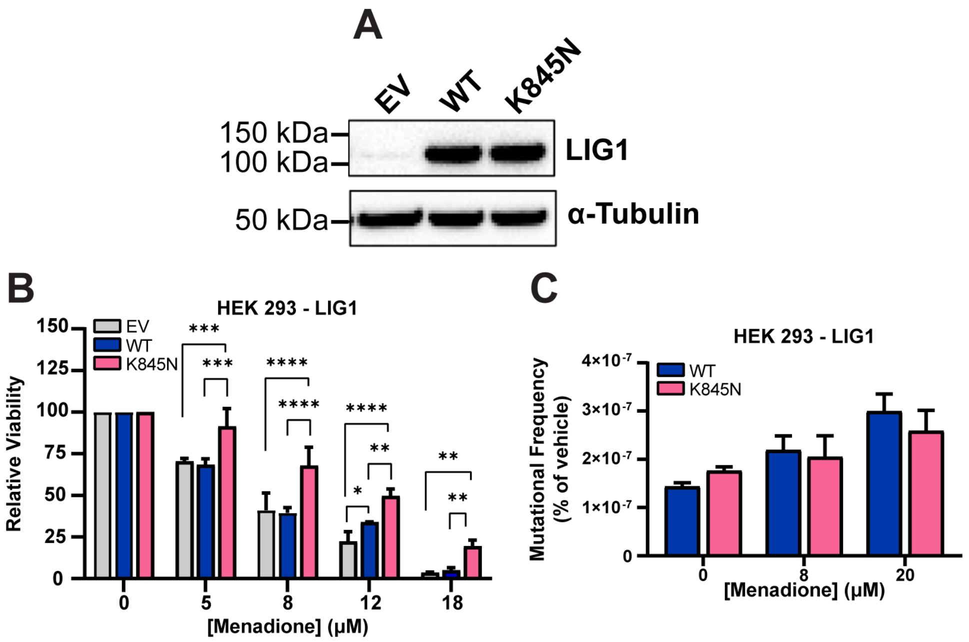


**Figure S7. Comparison of mutation frequency of HEK 293 cells expressing LIG1 WT or K845N under oxidative stress conditions**

**(**A) Expression of empty vehicle (EV), LIG1 WT and K845N in stably transfected HEK 293 cell lines was analyzed via western blot with anti-LIG1 and anti-alpha tubulin.

(B) HEK 293 cells (EV, WT or K845N) were treated with menadione for 4 hours at the indicated concentrations. Cells were washed in fresh medium and incubated for 48 hours. Cellular viability was measured by CTG. Bar graph shows mean % viability relative to baseline (0% menadione) of 3 technical replicates ± standard deviation. Statistical significance was determined from 2-way ANOVA with Tukey's multiple comparison test (****$p < 0.0001$, **$p < 0.01$, *$p < 0.05$). (C) HEK 293 cells (WT or K845N) were treated menadione for 4 hours at the indicated concentrations and washed in fresh medium. After 48 hours of incubation, cells were harvested for DNA extraction and mutation frequencies were determined by duplex sequencing. Bar graph shows mean mutation frequency (MF) of 2 technical replicates ± standard deviation.

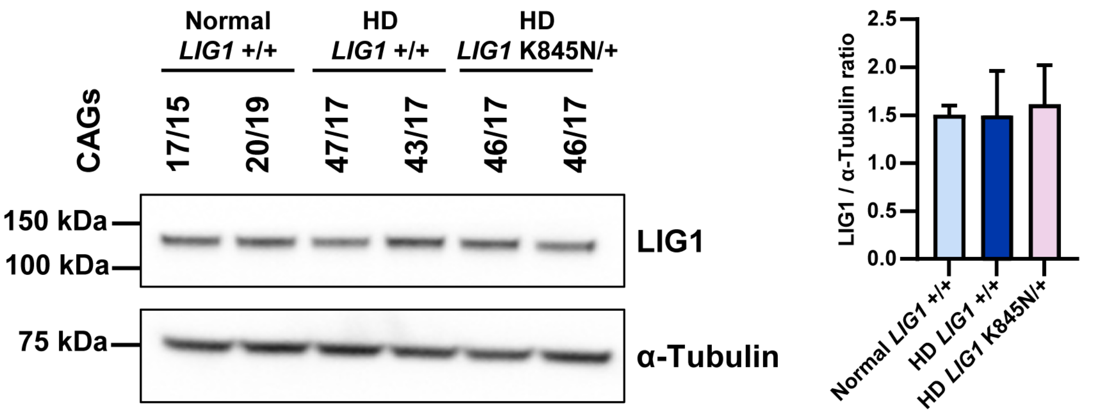


**Figure S8. The expression level of LIG1 in LCLs**

(A) The expression level of LIG1 was determined by western blot in normal and HD LCLs with anti-LIG1 and anti-alpha tubulin antibodies. *LIG1*+/+ denotes normal or HD LCLs homozygous for the *LIG1* rs145821638 (GRCh38 - Chr19:48117686) reference C-allele. *LIG1*K845N/+ denotes HD LCLs heterozygous for the 19AM3 modifier variant rs145821638 A-allele [17]. (B) The intensity of each band was quantified using image J, and LIG1 intensity normalized to alpha tubulin.

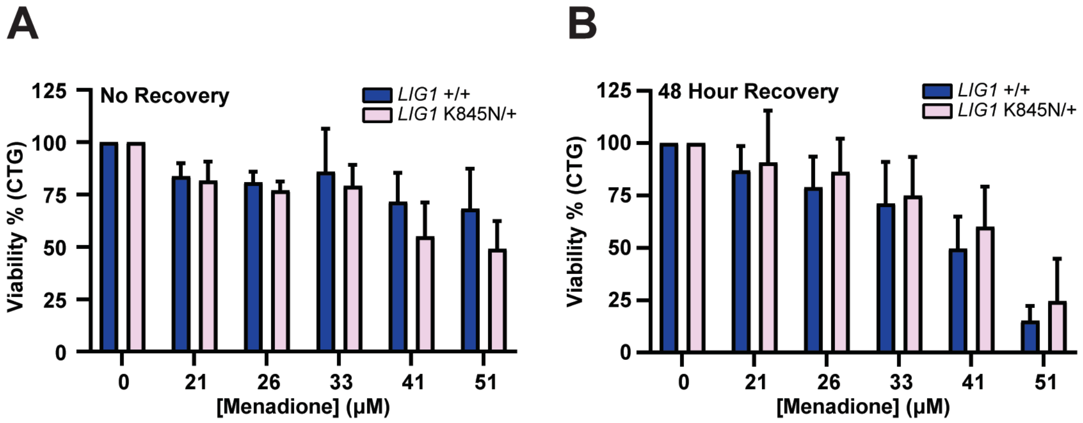


**Figure S9 Comparison of cytotoxicity of HD LCLs expressing LIG1 WT or K845N under menadione-induced stress conditions**

(A-B) HD LCLs homozygous for the *LIG1* rs145821638 (GRCh38 - Chr19:48117686) reference C-allele (*LIG1*+/+) or heterozygous for the 19AM3 modifier variant rs145821638 A-allele (*LIG1*K845N/+) [17] were treated with menadione for 4 hours at the indicated concentrations (see Fig.3 for details of LCLs). (A) Cell viability was measured immediately with no recovery time. (B) Cells were washed in fresh medium and incubated for 48 hours. Cellular viability was measured by CTG from the mean value of three technical replicates. Bar graph shows mean % viability relative to baseline (0% menadione) ± standard deviation (n = 4 *LIG1* +/+ independent LCLs, n = 3 independent *LIG1* K845N/+ LCLs). Statistical significance indicated in figure was determined by 2-way ANOVA with Tukey's multiple comparison test.

| *Lig1* genotype | *Htt* genotype | Males | Females | Total |
|---|---|---|---|---|
| K834N/K843N | Q111/+ | 11 | 6 | 17 |
| | +/+ | 7 | 16 | 23 |
| | Combined | 18 | 22 | 40 |
| K834N/+ | Q111/+ | 28 | 27 | 55 |
| | +/+ | 23 | 25 | 48 |
| | Combined | 51 | 52 | 103 |
| +/+ | Q111/+ | 5 | 16 | 21 |
| | +/+ | 12 | 14 | 26 |
| | Combined | 17 | 30 | 47 |

| Test | Chi2, df | P val |
|---|---|---|
| Males, Observed vs. Expected, Combined *Htt* genotypes | 1.716, 2 | 0.4240 |
| Females, Observed vs. Expected, Combined *Htt* genotypes | 0.6190, 2 | 0.7338 |
| Total, Observed vs. Expected,, Combined *Htt* genotypes | 1.058, 2 | 0.5891 |
| Total, *Htt*Q111/+ vs. *Htt*+/+ | 1.824, 2 | 0.4017 |

**Figure S10. Expected Mendelian ratios in transmission of the *Lig1*$^{K843N}$ allele.**

Top: Number of pups (genotyped at weaning) in *Htt*$^{Q111/+}$ *Lig1*$^{K843N/+}$ x *Htt*$^{+/+}$ *Lig1*$^{K843N/+}$ crosses. Bottom: Chi square tests show lack of deviation from expected Mendelian ratios and no difference in Mendelian ratios between *Htt*$^{+/+}$ and *Htt*$^{Q111/+}$ backgrounds.

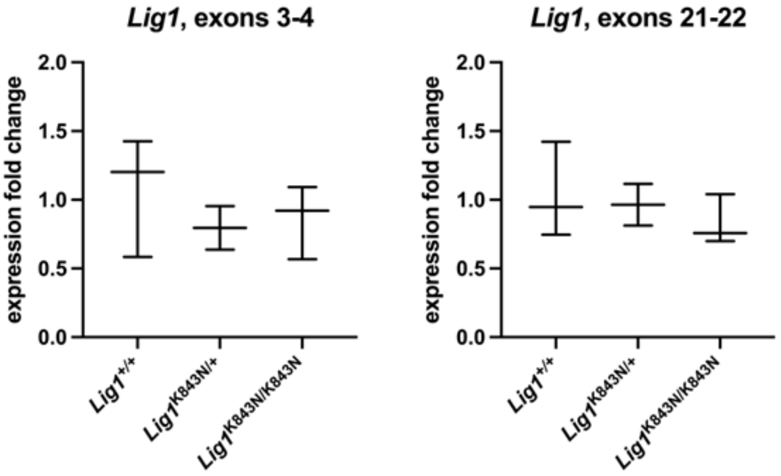


**Figure S11. Impact of *Lig1*$^{K843N}$ variant on *Lig1* expression**

Quantitative PCRs using TaqMan assays of *Lig1* mRNA (exons 21-22 and exons 3-4) in mouse liver. Relative expression values ($DC_t$) of *Lig1* for each mouse were calculated using the geometric mean of the $C_t$ values of three houskeeping genes *Ppia*, *Actb* and *Gusb*. Expression fold change ($2^{-DDCt}$) of *Lig1* for each mouse was calculated relative to the geomean of relative expression values ($DC_t$) of the control group (*Lig1*$^{+/+}$) mice. One way ANOVA with Tukey's multiple comparison test did not show significant differences between genotypes. *Lig1*$^{+/+}$ N=3, *Lig1*$^{K843N/+}$ N=2, *Lig1*$^{K843N/K843N}$ N=3. Two technical replicates were performed for each mouse tissue, which were averaged.

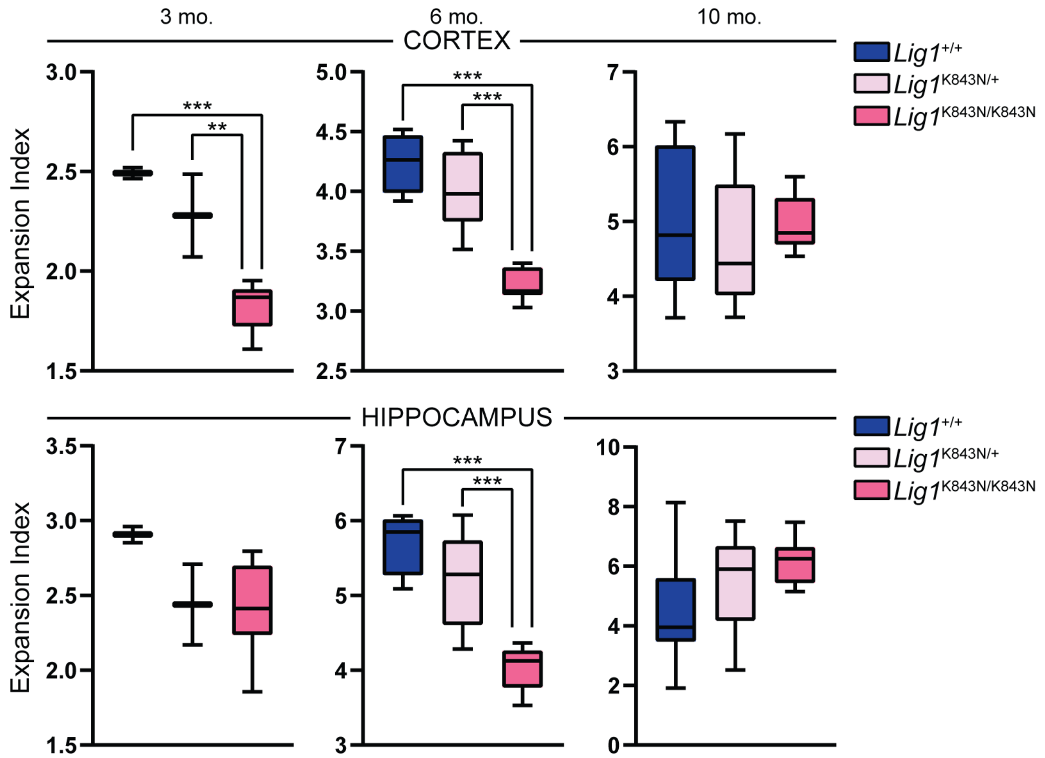


**Figure S12. Impact of *Lig1*$^{\text{K843N}}$ variant on somatic expansion in cortex and hippocampus**

Somatic CAG expansion indices (Min to Max box-whisker plots) from cortex and hippocampus of *Htt*$^{\text{Q111/+}}$ mice with different *Lig1* genotypes. 3 mo: *Lig1*$^{+/+}$ N=2, *Lig1*$^{\text{K843N/+}}$ N=2, *Lig1*$^{\text{K843N/K843N}}$ N=8. 6 mo: *Lig1*$^{+/+}$ N=4, *Lig1*$^{\text{K843N/+}}$ N=9, *Lig1*$^{\text{K843N/K843N}}$ N=7. 10 mo: Lig1+/+ N=9, *Lig1*$^{\text{K843N/+}}$ N=9, *Lig1*$^{\text{K843N/K843N}}$ N=10. **$p < 0.01$; ***$p < 0.001$, ****$p < 0.0001$ (One way ANOVA, comparing all genotypes for each age and tissue, with Tukey's multiple comparison test).

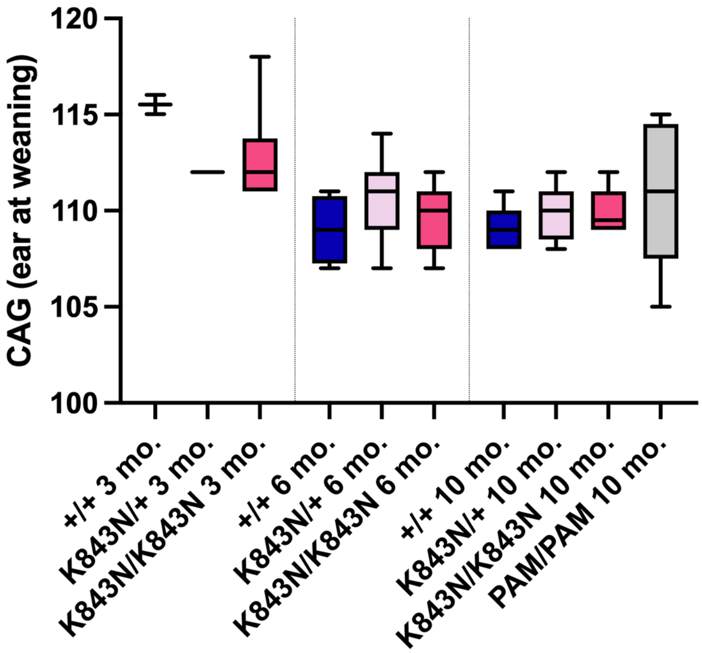


**Figure S13. Inherited CAG lengths in mice cohorts**

CAG lengths (Min to Max box-whisker plots) measured in ear tissue at weaning in 3-month, 6-month and 10-month cohorts. 3 mo: *Lig1*$^{+/+}$ N=2, *Lig1*$^{K843N/+}$ N=2, *Lig1*$^{K843N/K843N}$ N=8. 6 mo: *Lig1*$^{+/+}$ N=4, *Lig1*$^{K843N/+}$ N=9, *Lig1*$^{K843N/K843N}$ N=7. 10 mo: Lig1+/+ N=9, *Lig1*$^{K843N/+}$ N=9, *Lig1*$^{K843N/K843N}$ N=10, *Lig1*$^{PAM/PAM}$ N=5. *Lig1*$^{PAM/PAM}$ mice are homozygous for the silent C>T PAM mutation but are wild-type for the K843N variant. One way ANOVA, comparing all genotypes for each age, with Tukey's multiple comparison test, showed no significant differences in inherited CAG lengths between mice of different *Lig1* genotypes.

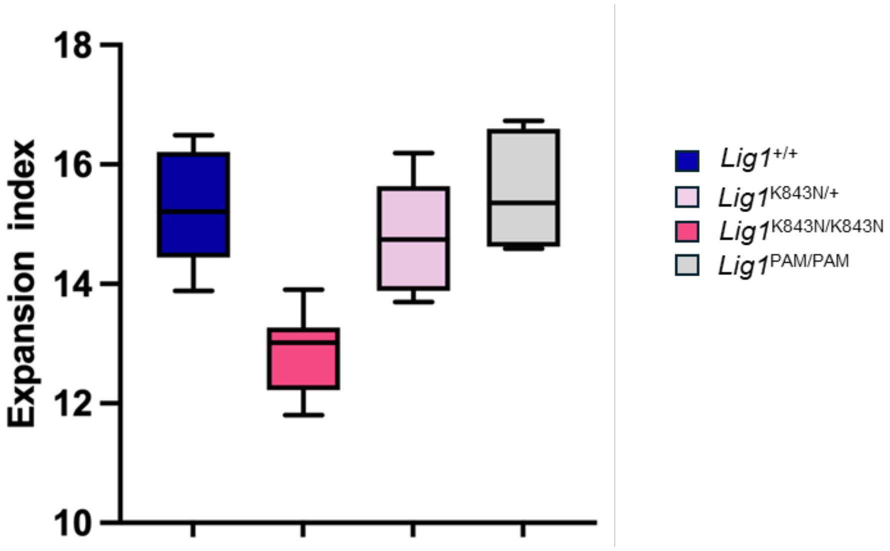


**Figure S14. Silent PAM mutation alone does not alter CAG expansion**

Somatic CAG expansion indices (Min to Max box-whisker plots) from liver of 10-month *Htt*$^{Q111/+}$ mice with different *Lig1* genotypes. *Lig1*$^{+/+}$ N=9, *Lig1*$^{K843N/+}$ N=9, *Lig1*$^{K843N/K843N}$ N=10, *Lig1*$^{PAM/PAM}$ N=5. One way ANOVA with Tukey's multiple comparison test comparing all genotypes, showed no significant difference between *Lig1*$^{+/+}$ and *Lig1*$^{PAM/PAM}$ expansion indices (p=0.9552), and significantly lower expansion indices in *Lig1*$^{K843N/K843N}$ mice compared to *Lig1*$^{PAM/PAM}$ mice ($p < 0.0001$). *Lig1*$^{PAM/PAM}$ mice are homozygous for the silent C>T PAM mutation but are wild-type for the K843N variant.

**Table S1. Oligonucleotide for ligation assay**

| Oligo Name | Sequence (5' – 3') |
|---|---|
| Up18OH-A | CAT GGG CGG CAT GAA CCA |
| Up18OH-G | CAT GGG CGG CAT GAA CCG |
| Up18OH-C | CAT GGG CGG CAT GAA CCC |
| Up18OH-T | CAT GGG CGG CAT GAA CCT |
| Up18OH-$^{oxo}$G | CAT GGG CGG CAT GAA CC$^{oxo}$G |
| DownP16-FAM | PO4-GAG GCC CAT CCT CAC C-FAM |
| Temp34-T | GGT GAG GAT GGG CCT CTG GTT CAT GCC GCC CAT G |
| Temp34-C | GGT GAG GAT GGG CCT CCG GTT CAT GCC GCC CAT G |
| Temp34-A | GGT GAG GAT GGG CCT CAG GTT CAT GCC GCC CAT G |
| Temp34-G | GGT GAG GAT GGG CCT CGG GTT CAT GCC GCC CAT G |
| Up13OH-C | GTGCTGATGCGTC |
| UP13OH-$^{oxo}$G | GTGCTGATGCGT$^{oxo}$G |
| DownP15-FAM | PO4-GTCGGACTGATTCGG-FAM |
| Temp28-G | CCGAATCAGTCCGACGACGCATCAGCAC |
| Temp28-A | CCGAATCAGTCCGACAACGCATCAGCAC |

The indicated Up18OH, P16-FAM, and Temp34 strand (Caglayan 2017) were annealed for ligation assays with the full-length LIG1. The indicated Up13OH, DownP15-FAM, and Temp28 strand (Tumbale 2019) were annealed for assays with Δ232 LIG1. Annealing ratios were 1:1.5:2 with respect to the phosphate, template, and 3′OH strands.

**Table S2. Steady-state kinetic parameters for Δ232 LIG1**

| | C:G Nicked DNA | | |
|---|---|---|---|
| | WT | K845N | Ratio K845N to WT |
| $k_{cat}$ ($s^{-1}$) | 0.52 ± 0.01 | 0.20 ± 0.01 | 0.38 |
| $K_M$ (nM) | 41.6 ± 5.9 | 80.7 ± 13.8 | 1.9 |
| $k_{cat}/K_M (M^{-1}s^{-1})$ | 12.5 ± 1.8 $x10^6$ | 2.5 ± 0.4 $x10^6$ | 0.20 |
| Abortive Ligation | 0.01 ± 0.00 | 0.028 ± 0.003 | |

| | 8oxoG:A Nicked DNA | | |
|---|---|---|---|
| | WT | K845N | Ratio K845N to WT |
| $k_{cat}$ ($s^{-1}$) | 0.08 ± 0.01 | $3x10^{-3}$ ± $5x10^{-4}$ | 0.038 |
| $K_M$ (nM) | 31.7 ± 5.9 | 73.3 ± 16.5 | 2.3 |
| $k_{cat}/K_M (M^{-1}s^{-1})$ | 2.4 ± 0.5 $x10^6$ | 0.04 ± 0.01 $x10^6$ | 0.017 |
| Abortive Ligation | 0.51 ± 0.03 | 0.93 ± 0.01 | |

| | Fidelity Measurements | | |
|---|---|---|---|
| | WT | K845N | Ratio K845N to WT |
| Discrimination against 8oxoG:A | 5.1 ± 1.3 | 60.1 ± 19.6 | 12 |

These data are from experiments conducted in Fig. 2. Reactions were performed in standard reaction buffer at 1.2 mM $MgCl_2$ (1 mM Free $Mg^{2+}$) and 0.2 mM ATP. Values are the average ± SD (n ≥ 3).
